# A robust and versatile nanobody platform for drug delivery

**DOI:** 10.1101/2020.08.19.257725

**Authors:** Zhuolun Shen, Yufei Xiang, Sandra Vegara, Apeng Chen, Zhengyun Xiao, Ulises Santiago, Changzhong Jin, Zhe Sang, Jiadi Luo, Kong Chen, Dina Schneidman-Duhovny, Carlos Camacho, Guillermo Calero, Baoli Hu, Yi Shi

## Abstract

Therapeutic and diagnostic efficacies of numerous small biomolecules and chemical compounds are hampered by the short half-lives. Here we report the development of a repertoire of diverse, high-affinity albumin-nanobodies (Nb_HSA_) to facilitate drug delivery. By integrating biophysics, and hybrid structural biology, we have systematically characterized the Nb_HSA_ for albumin binding, mapped the epitopes, and resolved the architecture of a tetrameric Nb-albumin complex. We employed quantitative proteomics for accurate, multiplex Nb pharmacokinetic analysis. Using a humanized albumin mouse model, we found that the Nb_HSA_ has outstanding pharmacokinetics; the most stable Nb_HSA_ has a 771-fold T_1/2_ improvement compared with a control Nb. Interestingly, the pharmacokinetics of Nb_HSA_ is related to their biophysical and structural properties. To demonstrate the utility of Nb_HSA_, we developed a highly stable Nb_HSA_-hIL-2 cytokine conjugate “Duraleukin” and confirmed its improved anticancer properties than hIL-2 alone. We envision that this high-quality Nb resource will advance research into novel biotherapeutics.

## Introduction

Numerous chemical compounds and small biologics, such as hormones, cytokines, and coagulation factors, have limited therapeutic efficacy due to their poor *in vivo* stabilities. Different strategies have been implemented to address this challenge, including chemical modifications, conjugation with the antibody Fc domain, and targeting to human serum albumin (HSA). Human serum albumin (HSA) is an abundant serum protein with an exceptionally long half-life of ~ three weeks ^1^. Unlike small biomolecules (< 50 kDa) rapidly eliminated (e.g., from mins to a few hours) *via* glomerular filtration, HSA can be retained due to the relatively large size (66.5 kDa). Importantly, HSA can bind with the neonatal Fc receptor (FcRn) at acidic pH and is continuously recycled *via* the FcRn-mediated transcytosis ^2,3^. Because of its stable interaction with the FcRn, it can escape the endo-lysosomal degradation pathway and obtain outstanding serum stability. Interestingly, although HSA is highly abundant in the serum, its intracellular concentration is likely very low ^3,4^.

The high *in vivo* solubility and stability of HSA make it an attractive endogenous carrier for drug delivery. Different HSA-based strategies, such as direct fusion to HSA, albumin-binding domains (ABDs) derived from gram-positive bacteria, and albumin nanoparticles, have been developed ^1,3,5^. Despite the encouraging progress, however, successful applications of these methods towards robust bioengineering and clinical uses remain limited ^6^. The development of robust agents, ideally easily accessible to the research community, will help advance drug development.

Nanobodies (Nbs) are small antigen-binding fragments (~15 kDa) derived from the camelid’s heavy chain only antibodies (HcAbs) ^7^. They are well known for the robust folds, excellent solubility, stability, and ease of bioengineering and manufacturing. Besides, they can efficiently penetrate tissues and are potentially low immunogenic for clinical applications due to the high sequence similarity with the human IgGs. Because of these characteristics, Nbs have emerged as a new class of antibodies for biomedical research and drug development ^8,9^.

We have recently discovered an extensive repertoire of anti-HSA Nbs (Nb_HSA_) by camelid immunization of recombinant HSA in conjugation to proteomic identifications {*Xiang* et al., submitted}^10^. We selectively produced a set of diverse Nb_HSA_ sequences for systematic characterizations by multidisciplinary approaches, including biophysics, integrated proteomics, structural biology, and modeling. We then developed a strategy based on PRM (parallel reaction monitoring) mass spectrometry that enabled unbiased, parallel pharmacokinetic (PK) analysis of dozens of high-affinity Nb_HSA_ in a humanized albumin mouse model. Our studies revealed that the Nb_HSA_ PKs are related to their binding kinetics at the acidic pH environment and their epitopes. To demonstrate the utility of our Nb_HSA_, we conjugated high-affinity, cross-species Nbs to a critical cytokine interleukin-2 (hIL-2), developed and characterized several stable Nb-cytokine conjugates (Duraleukins), and have confirmed the improved antitumor efficacy of a Duraleukin than hIL-2 in a melanoma mouse model. We envision that this high-quality Nb resource and the hybrid proteomic technologies will find broad utility in biomedical applications.

## Results

### Characterization of a highly diverse repertoire of high-quality Nb_HSA_

A collection of 71 recombinant His6-tagged Nbs with diverse sequences was produced from *E. coli* cells (**Fig 1a, Table S1**). The selected Nb_HSA_ vary significantly on the CDR3 (complementarity-determining region 3) fingerprint regions, including amino acid compositions, sequence length, isoelectric points (pIs), and hydrophobicity potentially spanning a large spectrum of physicochemical structural properties (**Fig 1b**). We performed enzyme-linked immunosorbent assay (ELISA) to assess their relative affinities and cross-species selectivities. We ranked Nb_HSA_ based on the ELISA affinities to different albumins, including HSA, cynomolgus monkey albumin, and mouse albumin. HSA is highly conserved with cynomolgus monkey albumin (92% sequence identity) and is relatively conserved with mouse albumin (78% sequence identity). As expected, greater than 75% (53 Nb_HSA_) of Nb_HSA_ can bind monkey albumin, and 11% (8 Nb_HSA_) can bind mouse albumin (**Fig 1c, Methods**). Interestingly, 8% (6 Nb_HSA_) strongly associated with all three albumin orthologs and were confirmed by immunoprecipitations (**Fig 1c**). For example, Nb_132_ binds both humans, monkeys, and mouse albumin but not bovine and llama albumin (**Fig 1g**). The high affinity, cross-species Nb_HSA_ are particularly useful for Nb-based drug development in the animal models.

**Figure 1.**
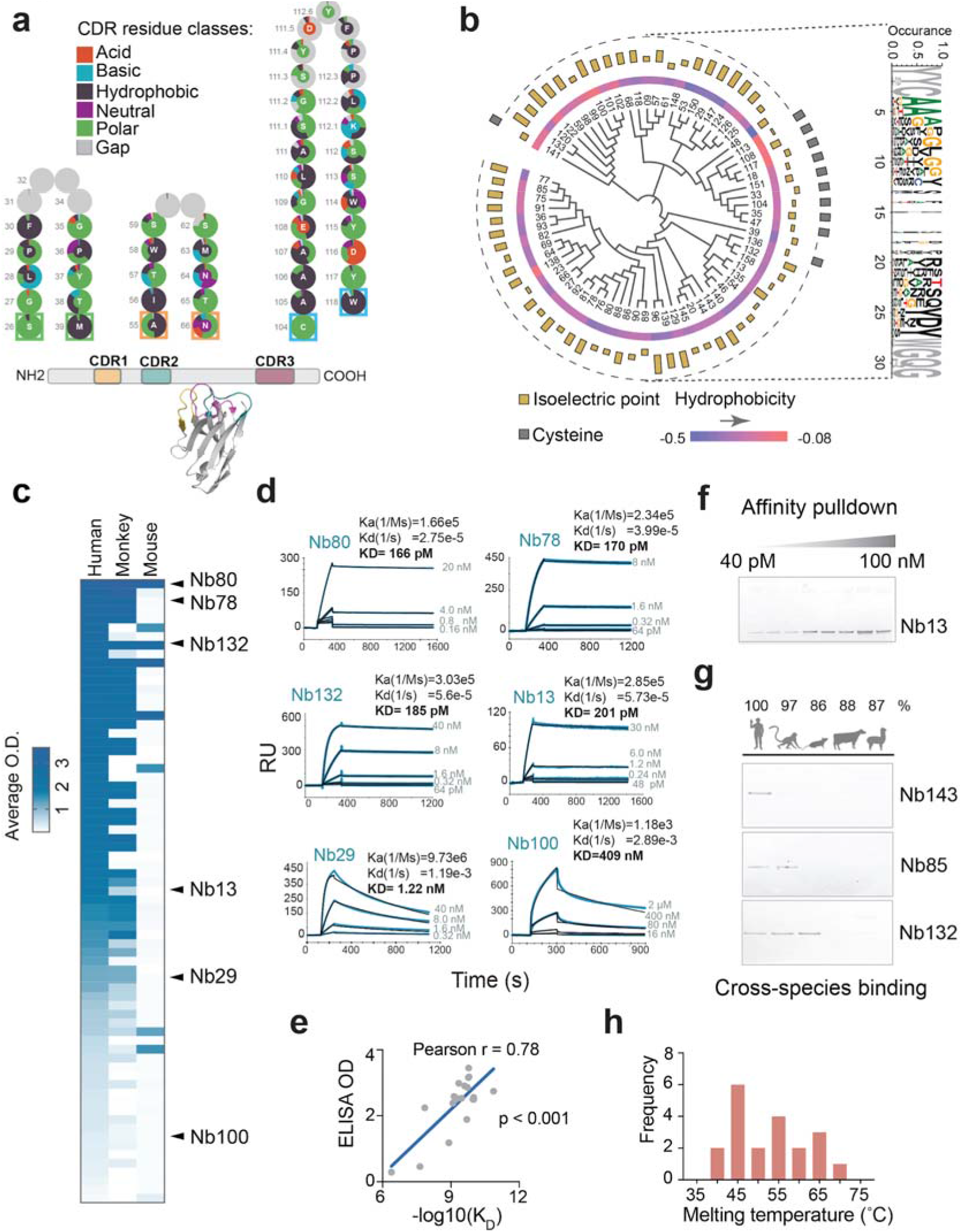
Identification and characterization of Nb_HSA_. a) Schematic structure and amino acid composition of Nb_HSA_. 71 different Nbs were analyzed by Yvis. The amino acid frequency at each CDR position was calculated. Amino acids were color-coded and classified based on the physicochemical properties. b) Circos and sequence logo plots showing the diversity of CDR3. c) ELISA heat map of albumin cross-species binding of 71 different Nbs. d) Binding kinetics of 6 representative Nbs by SPR (surface plasmon resonance). e) Correlation between ELISA O.D. (optical density) and SPR K_D_ affinity. f) Immunoprecipitation of the HSA-Nb_13_ complex at different concentrations of Nb_13._ g) Validation of Nb cross-species reactivity by immunoprecipitation. Three Nbs (Nb_143_, Nb_85_, and Nb_132_) were immunoprecipitated by affinity resins coupled with albumin of different species, including human, monkey, mouse, bovine, and llama. h) The plot of the thermostability of 20 Nb_HSA_ selected for measurement by differential scanning fluorimetry.

We selected 20 Nb_HSA_ of high yields (from *E. coli* cell lysis) to assess the binding kinetics by surface plasmon resonance (SPR) (**Fig 1d, Fig S1**). We found a good correlation (Pearson r= 0.78, p<0.001) between the ELISA affinity and SPR K_D_ (**Fig 1e**). 75% (15 out of 20) of Nb_HSA_ present sub-nM K_D_ with different binding kinetics, while the remaining 25% have single-digit to hundreds of nM affinities (**Fig 1f**). Next, we evaluated the thermostability of Nb_HSA_ by differential scanning fluorimetry. The Nbs are generally thermostable, with melting temperatures between 39 □C to 70 □C (**Fig 1h**, **Fig S2, Table S2**). The excellent physicochemical properties, including solubility, target specificity, affinity, and stability, are essential for Nb therapeutic development ^11,12^.

### Integrative structural proteomics reveals dominant epitopes for Nb_HSA_ binding

Epitope mapping provides insight into antigen-antibody interactions and is useful for rational drug designs ^13–15^. We employed an integrative structure approach by chemical cross-linking/mass spectrometry (CXMS) and computational modeling to map HSA epitopes for Nb binding. CXMS identifies residue-specific cross-link restraints that are highly informative to interrogate the binding interfaces of protein complexes. Cross-linking information can be readily integrated into other structural biology techniques and computational tools to derive accurate structural models ^16–21^.

We purified and reconstituted 38 different HSA-Nb complexes for disuccinimidyl subtracted (DSS) crosslinking, which can reach up to 30 Å between two cross-linked amines (e.g., lysine residues). A total of 125 intersubunit cross-links were confidently identified by high-resolution MS (**Table S3, Methods**). These cross-links were then used to compute structural models. Integrative modeling analysis revealed that 29 HSA-Nb complexes converged with an average RMSD (root-mean-square-deviation) of 7.4 Å (**Fig 2c**). Overall, 87.5% of cross-links were satisfied by the models (i.e., <30 Å) (**Fig S3b**).

**Figure 2.**
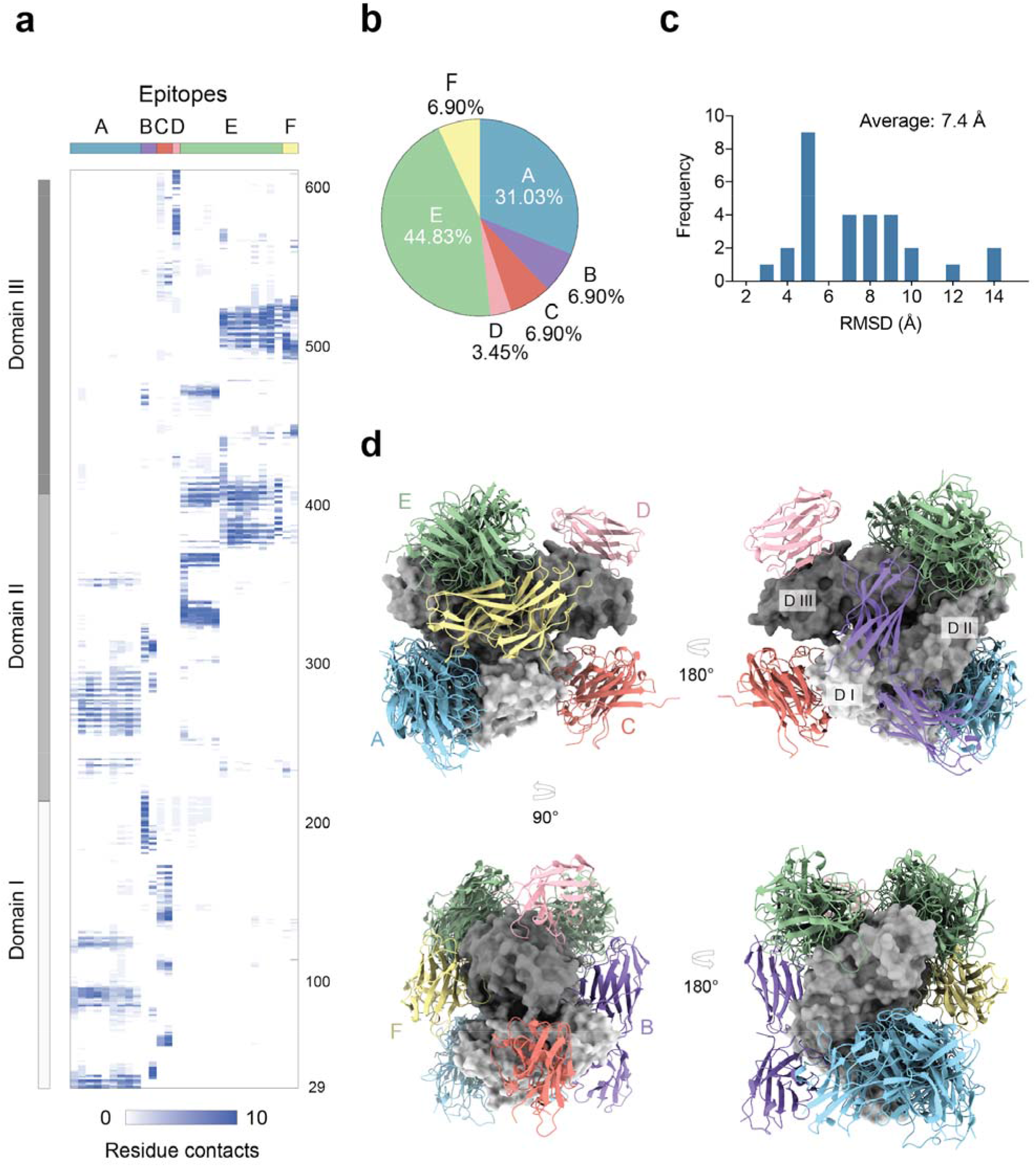
Cross-link models of the HSA-Nb_HSA_ complexes. a) The epitope clusters of HSA based on converged cross-link models. HSA Epitope cluster A: residues 29-37, 81-93, 118-124, 232, 236, 252-290, 300-310, 345-350; cluster B: residues 35-44, 179-212, 282-286, 298-318, 460-465; cluster C: residues 56-62, 101-108, 1234-169; cluster D: residues 426, 566-582, 595, 598-606; cluster F: 440-446, 488-525. Cluster E has two subtypes that share the same helices region and vary on other binding residues. E_type1_: residues 321-331, 334-335, 358, 360-365, 395-413, 465-470; E_type2_: residues 371-387, 396-410, 494-519. b) Percentage of epitope-specific Nb_HSA_ based on converged cross-link models. c) The RMSD (root-mean-square deviation, Å) distribution of the cross-link models. d) Integrative structural models of HSA-Nb_HSA_ complexes. Only the top-scored Nb_HSA_ models were shown. Different molecules were presented in different colors. Grey (surface presentation): HSA (with three domains DI, DII, and DIII). Nb_HSA_ corresponding to different epitopes were presented as cartoons. Epitope Cluster A: blue; B: purple; C: salmon; D: light pink; E: green; F: yellow.

Cross-linking based structural modeling identified 6 HSA epitope clusters (from A to F) of Nb_HSA_ (**Fig 2a, Table S3**). Despite the high sequence diversity of CDR3s, >75% of the Nb_HSA_ (21 Nbs) binds two dominant epitope clusters: clusters A (31.03%) and E (44.83%) (**Fig 2b**), indicating a convergent, CDR3-dependent contact between Nbs and HSA. We found excellent shape complementarities between the concave epitopes on HSA and the corresponding convex paratopes of Nb_HSA_ for high-affinity interactions (**Fig 2d**). Interestingly, a sub-nM affinity Nb (Nb_126_) was identified to localize at the major crevice on HSA that regulates FcRn binding (**Fig S3a**), raising a possibility of competitive binding between Nb126 and the FcRn.

### Hybrid structural determination of a tetrameric HSA-Nb complex

To further verify our integrative structures, we selected Nb_13_, Nb_29_, and Nb_80_ that correspond to three epitope clusters A, C, and E, respectively, and employed size-exclusion chromatography (SEC) to confirm that they do not compete for HSA binding. The presence of three separate SEC peaks corresponding to HSA-Nb complexes of different molecular weights indicates that these Nbs bind distinct HSA epitopes (**Fig 3a**). After SEC, a homogenous fraction of the complex was selected for analysis by negative stain electron microscopy (EM). ~22,000 EM particles were selected to reconstruct a 3D density map (**Fig 3b, Methods**). Consistently, the cross-link models of the tetrameric Nb-HSA complex agreed well with EM density (**Fig 3c**). Based on our integrative structure, we found that sub-nM Nb_80_ (K_D_=166 pM) and Nb_13_ (K_D_=201 pM) have better shape complementarity with HSA than Nb_29_ (K_D_=1.2 nM) (**Fig 3d-f**). Our models revealed the formation of two crossed, salt bridge pairs between HSA_K383_ and HSA_E400_ with the respective CDR residues on Nb_80_. To validate this, we made two HSA mutants and evaluated their binding to Nb_HSA_ (**Fig 3g**). While a single-point mutation E400R significantly weakened the HSA-Nb_80_ association, a double-residue mutation (K383D-E400R) completely abolished the strong interaction (**Fig 3h**). Collectively, our integrative structure of the tetrameric HSA-Nb complex confirmed the epitope mapping results and provided the structural basis to understand the HSA-Nb interactions.

**Figure 3.**
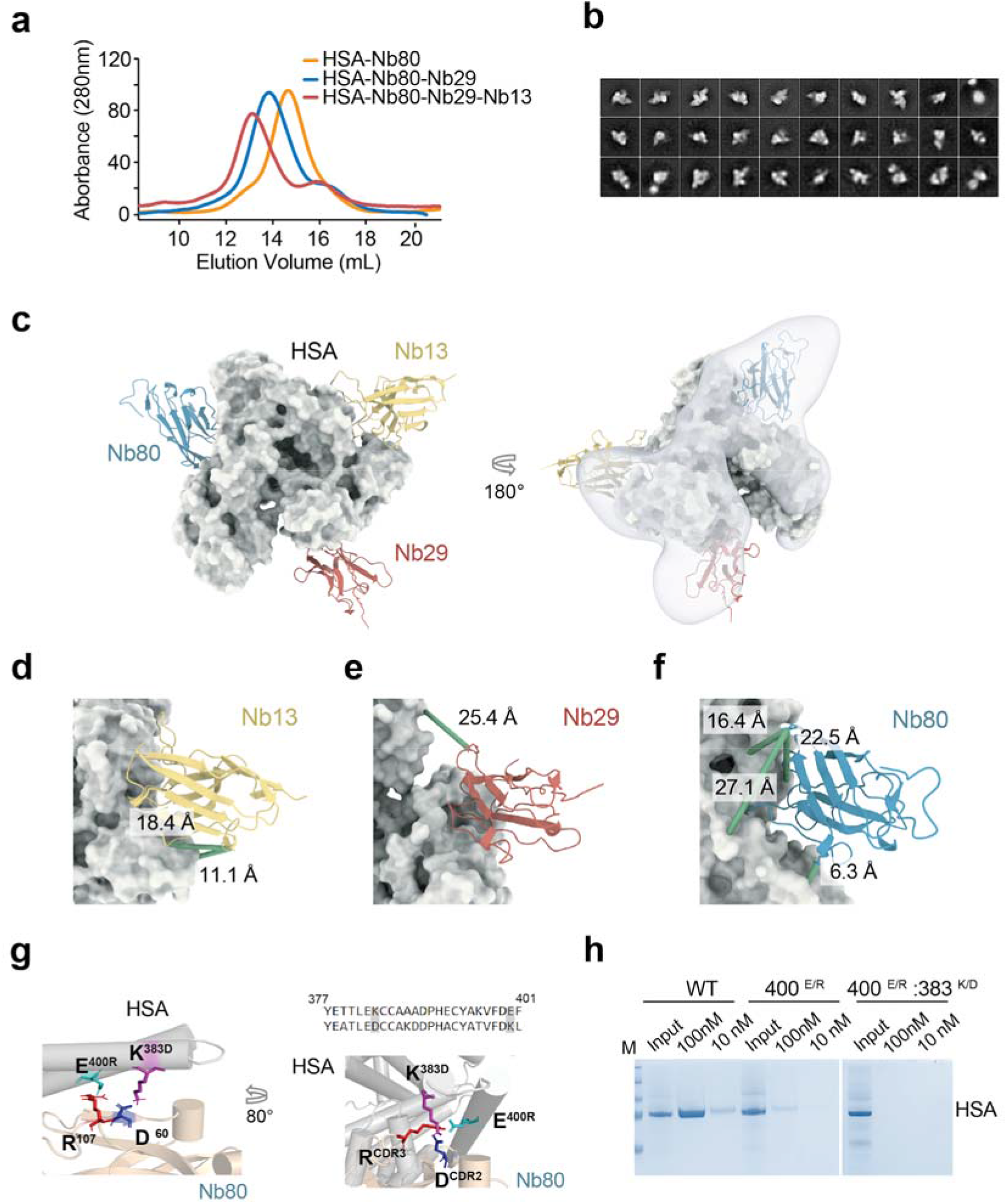
Integrative structural characterization of a tetrameric HSA-Nb complex. a) Size-exclusion chromatography (SEC) analysis of the reconstituted tetrameric complex, including Nb_13_, Nb_29_, Nb_80_, and HSA. b) Representative negative stain EM images of the tetrameric complex. c) Overlapping of the cross-link structure model of the tetrameric complex with the negative stain EM structure. Nb_HSA_ were presented in different colors. Yellow: Nb13; salmon: Nb29; blue: Nb80. d)-f) Close-up views of the interfaces of the tetrameric HSA-Nb complex and the cross-link restraints. g) A cross-link model of HSA and Nb80 interaction showing the putative salt bridges. Sidechains of two charged residues on HSA (K383 and E400) and the corresponding residues on Nb80 were shown. The HSA sequence (from residue 377 to residue 401) was aligned with camelid albumin. h) HSA site-directed mutagenesis and immunoprecipitation of the mutant HSA. Nb_80_-conjugated resins were used to pull down different HSA mutants, including E400R and E400R-K383D.

### Development of a proteomic approach that enables specific, accurate and multiplex Nb pharmacokinetic analysis

ELISA has been extensively used for the analysis of antibody PK. However, this approach can be significantly limited by the availability of high-quality detection antibodies, rendering specific and accurate PK analysis from the complex serum background challenging. Such a method is particularly challenging for comparative PK analysis of a large set of antibodies as it requires a substantial cohort of animals and slow handling time that may complicate experiments and introduce bias. To alleviate these problems, here we developed a strategy by employing MS2-based quantitative proteomics (parallel reaction monitoring or PRM) to facilitate specific, accurate, and multiplex analysis of Nb PKs from the serum samples. This approach enables specific detections of at least three femtomoles of an Nb (10^−15^ mole) per μL of the serum and offers better quantification linearity than the MS-1 based approach (R^2^= 0.999) (**Fig S5, Table S4**). The high dynamic range of this approach is essential for accurate PK measurement.

Since most of our Nb_HSA_ do not strongly bind mouse albumin, we chose a humanized albumin mouse model to mimic the physiology of endogenous HSA in humans. This mouse model (B6.Cg-Tg(FCGRT)32Dcr *Alb*^*em12Mvw*^ *Fcgrt*^*tm1Dcr*^/MvwJ) removed the native mouse albumin and replaced the mouse FcRn with a human ortholog. HSA can be later administered to study PK of HSA-based drugs ^22^.

Following HSA injection, we administered a mixture of 22 Nbs, including 20 well-characterized Nb_HSA_ (with SPR kinetic and thermostability measurements) and two non-binder control Nbs with equal molarity to this model (n = 3) *via* single bolus intravenous (*i.v.*) injections (**Methods**). We used a molar ratio of 1: 5 between Nb_HSA_ and HSA to ensure an excessive HSA, avoiding potential competitive binding among different Nb_HSA_. The blood was sampled at different time points post-injection. Sera fractions containing Nb_HSA_ and other serum proteins were efficiently proteolyzed. The resulting signature Nb_HSA_ peptides and their fragment ions were accurately quantified by PRM ^23,24^ (**Fig 4a, Table S4**). The median CV (coefficient of variation) of our proteomic quantifications from three animals was 15.3% (**Fig S6**).

**Figure 4.**
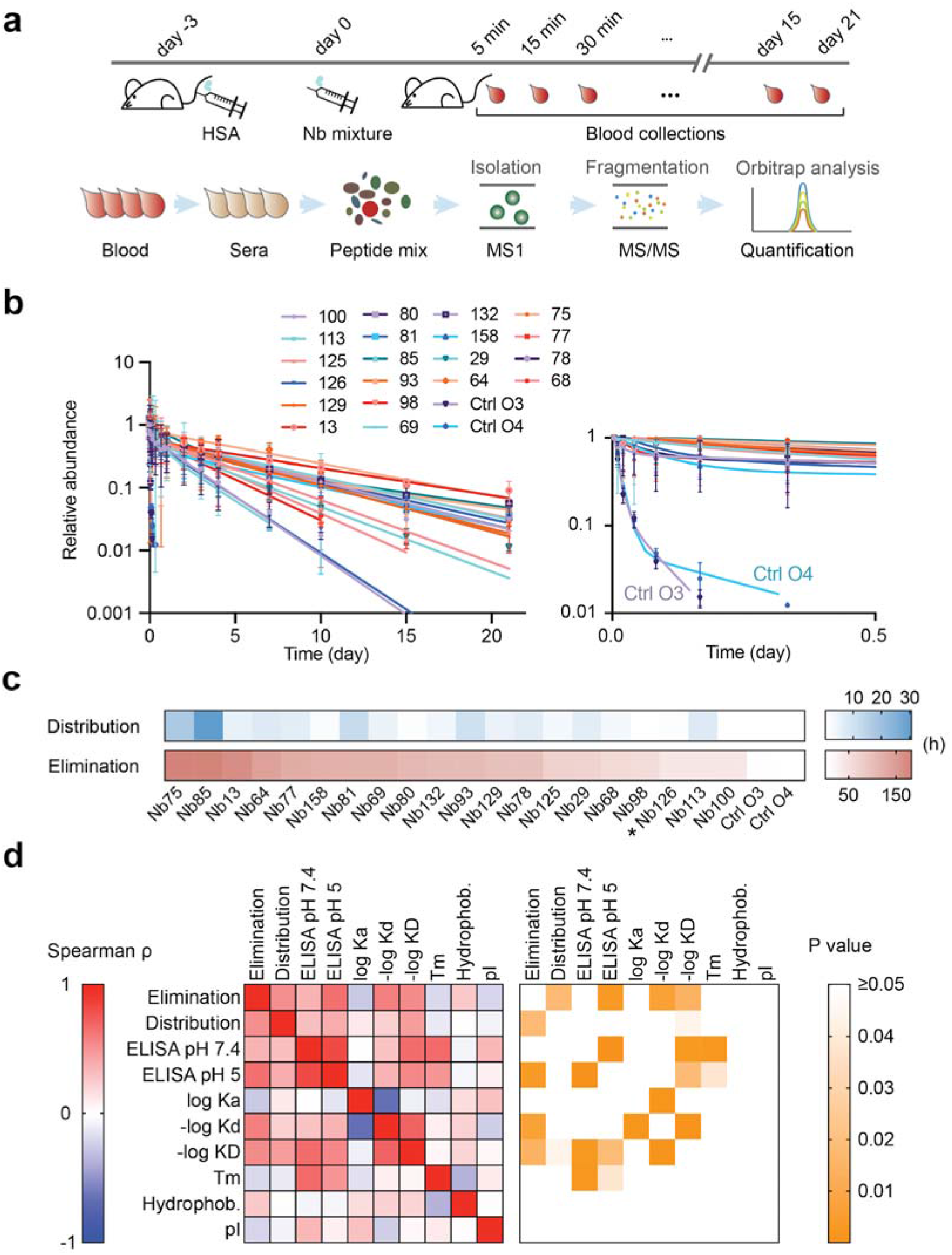
High-throughput Nb pharmacokinetics in a humanized albumin mouse model. a) Schematics of the MS-based assay for multiplexed pharmacokinetic (PK) measurement. Briefly, after consecutive injections of HSA and the Nb mixtures into the mouse model, blood was collected at 21 different time points and processed for proteomics-based pharmacokinetic analysis. b) PK analysis of 22 Nbs in a humanized albumin mouse model. Single bolus administration of an equimolar mixture of 22 Nbs, including 20 Nb_HSA_ and two non-binder controls were administered to three animals via *i.v.*injection. Serum samples were collected at different time points and proteolyzed. The resulting peptides were analyzed by LC coupled to MS (PRM). Each data point indicates the median Nb abundance from three different animals. The analysis was repeated three times. The data was then fitted into a two-phase decay model to calculate the Nb half-lives. c) Heatmap analysis of the distribution and elimination of Nb PK. *, See Fig S3A. d) Correlative analysis between PK and the properties of Nbs. For each correlation analysis, the correlation coefficient Spearman ρ, and the corresponding p-value were calculated using Prism GraphPad.

We observed that the control Nbs were rapidly cleared from the serum with the half-lives (T_1/2_s) of approximately 26-60 mins, consistent with the previous ELISA measurements ^25^. In sharp contrast, our Nb_HSA_ showed significantly increased T_1/2_s of up to 771-fold, from 1.6 days (Nb_100_) to 7.6 days (Nb_75_) (**Fig 4b, 4c, Table S2**). The T_1/2_s of Nb_HSA_ correlated with T_1/2_ of HSA (~ 8 days) that we measured in this model (**Fig S7c**). To explore the potential mechanism(s) that underlies the relatively fast T_1/2_ of HSA in this mouse model (the T_1/2_ of HSA in humans is ~21 days), we employed label-free quantitative proteomics ^26,27^ to monitor the longitudinal changes of the blood proteome of a mouse (up to 10 days) post-HSA injection. Four hundred forty-one blood proteins were accurately quantified and clustered based on their relative abundance changes (**Fig S8**). We found that a group of acute-phase inflammatory proteins, such as Serum Amyloid A and C-reactive protein, increased by hundreds of folds in abundance after 12-24 hours post-injection, before returning to the baseline level after 24-48 hours ^28,29^. Interestingly, we observed an increase of serum IgGs after 5-7 days indicating the potential anti-HSA immune response, which may explain the faster-than-expected clearance of the injected HSA in this model ^30^.

### Nb_HSA_ pharmacokinetics are related to their physicochemical and structural properties

The findings that Nb_HSA_ has diverse PK prompted us to investigate the potential mechanism(s) contributing to such differences. We analyzed the correlations between the PK and the physicochemical properties of Nb_HSA_, including affinity (at both neutral and acidic pHs), T_M_, pI, and hydrophobicity (**Fig 4d**). We found that Nb_HSA_ ELISA affinities at pH 5 (acidic pH) and SPR dissociation rates K_d_, positively correlated with the elimination rates (p < 0.01). In general, the higher the affinities, the slower Nb_HSA_ was cleared from the blood. It is conceivable that both the slow dissociation-rates and high-affinity binding at the acidic pH are important for the extended T_1/2_ of the low abundance Nb_HSA_-HSA-FcRn ternary complexes in the endosome. Nb_HSA_ with lower affinities at the acidic pH tend to dissociate faster from the complex and, therefore, have decreased T_1/2_. The only exception was Nb_126_, which binds strongly with HSA (K_D_ of 251 pM) and has relatively fast T_1/2_s (41.4 hrs). The interaction with HSA might be interfered with by a structural change of the epitope induced by FcRn binding (**Fig S3a**) ^2,30^.

### Development of Duraleukin - a novel application of the Nb_HSA_ platform for cancer immunotherapy

To demonstrate the utility of our Nbs for drug development, we applied them to modify a key cytokine interleukin-2 (IL-2). IL-2 functions primarily to regulate immune cells such as T cells and natural killer cells, thereby regulating immune responses that suppressor cancer ^31^. Human IL-2 (hIL-2) was the first FDA-approved cancer immunotherapy drug (in 1992) developed for effective treatment of renal cell carcinoma and metastatic melanoma ^32^. ~10% of these devastating late-stage cancer patients who received high dose hIL-2 treatment had a complete response. However, the clinical efficacy of hIL-2 has been significantly limited by its small size (~15 kDa) and poor PK (i.e., shorter than 30 minutes in humans) ^31,32^. High-dose, repetitive administration of hIL-2 (e.g., *i.v.* treatment every 8 hours), while critical for its antitumor efficacy, can lead to severe side effects such as liver toxicity and vascular leak syndrome ^32,33^.

We fused our Nb_HSA_ with hIL-2 to develop “Duraleukin” for mitigation of the poor PK of hIL-2 (**Fig 5a**). We chose three high-affinity Nb_HSA_ (Nb_77_, Nb_80_ and Nb_158_) that bind both human and mouse albumin to demonstrate the robustness of this approach. Large quantities of Duraleukins (DL_77_, DL_80_and DL_158_) were rapidly produced from *E. coli* (**Fig 5b**, **Methods**). All three DLs demonstrated excellent thermostability (T_m_ ≈ 60 °C, **Fig 5c**). They had comparable bioactivities to hIL-2 alone in the *in vitro* CD8^+^ T cell proliferation assay (**Fig 5d**), and were stable for days when incubated with mouse serum at 37°C (**Fig 5e**). Moreover, these DLs retained sub-nM affinities for HSA binding (**Fig 5e**). We selected DL_80_ for further *in vivo* evaluation because it has a high affinity for both mouse and human albumin binding, and remained associated with HSA at acidic pH (**Fig 5g**).

**Figure 5.**
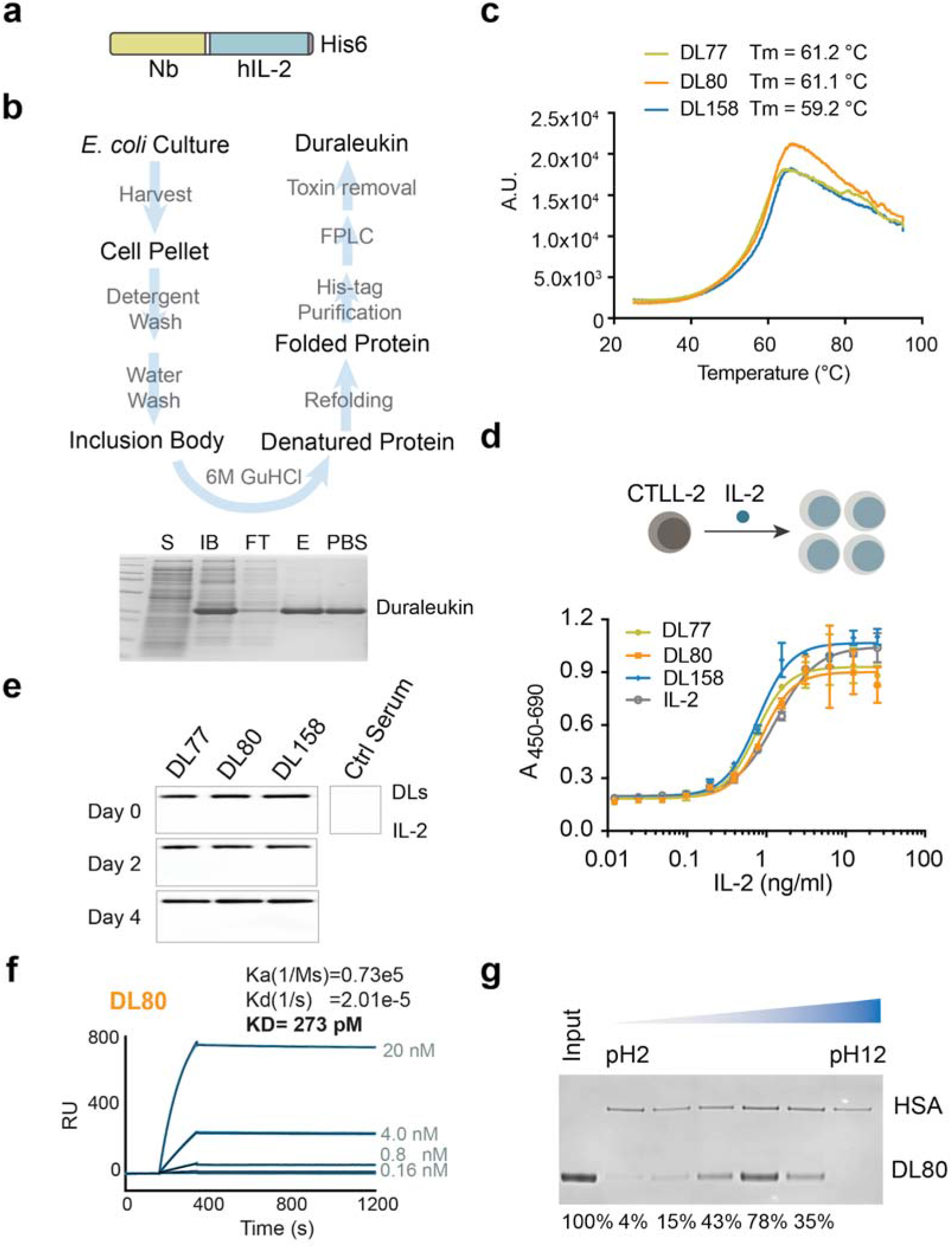
Development of Duraleukin- a novel class of Nb_HSA_-fusion cytokine. a) Schematic design of Duraleukin, including an N-terminal Nb_HSA_, a short flexible linker sequence ((GGGGS)_2_), and hIL-2 followed by a C-terminal His6 tag. b) Schematics of large-scale Duraleukin production from *E. coli*. A representative SDS-PAGE analysis of purified Duraleukin. S: soluble fraction. IB: inclusion body; FT: flow-through; E: elution; PBS: after endotoxin removal and dialysis in PBS. c) Thermostability of Duraleukins by differential scanning fluorimetry. d) *In vitro* CTLL-2 cell proliferation assay of Duraleukin and hIL-2. The X-axis is the concentration of hIL-2 (ng/ml). The Y-axis is the absorbance at 450 nM subtracted by absorbance at 690 nm (background signal) for the measurement of T cell proliferation. e) The resistance of three DLs to serum proteases. Purified DLs were incubated with the mouse serum for the indicated periods. After incubation, DLs were detected by western blot using an anti-IL-2 monoclonal antibody (BG5). f) The SPR kinetic measurement of DL80 for HSA binding. g) Immunoprecipitation assay (pH-dependent) of DL80 for HSA binding. HSA-conjugated agarose resin was used to pull down DL80 at various concentrations. Immunoprecipitated DL80 at different pH was normalized and quantified by ImageJ.

### Duraleukin demonstrates superior therapeutic efficacy for melanoma treatment in a mouse model

We confirmed that DL_80_ had a 46-fold improvement of PK compared with hIL-2 in C57BL/6J mouse (**Fig S9**). This relatively moderate improvement corresponds to the half-life of endogenous mouse albumin (<1 day) ^25^. We then assessed its therapeutic efficacy in the B16F10 melanoma model ^34^. Tumor-bearing C57BL/6J mice were intraperitoneally administered with equimolar of either DL_80_ or hIL-2 at each injection. The animals were treated with different frequencies (every six days for DL_80_ and hIL-2 daily) for a 24-day treatment. TA99 monoclonal antibody that recognizes melanoma marker TRP1 and functions synergistically with hIL-2, was co-treated for both groups ^35–38^ (**Methods**). We continuously monitored tumor sizes and survival for 62 days starting from the first treatment (**Fig 6a, 6b**). We observed that both DL_80_ and hIL-2 treated groups exhibited significant reductions in tumor loads and better overall survival than the PBS controls. As little as 1/6 total therapeutic dose was applied to the DL_80_-treated animals, and due to the relatively short T1/2 of mouse albumin, only a moderate PK improvement was observed in the C57BL/6J mice. Despite this, we observed near-complete tumor recessions for the DL_80_ responsive animals. In contrast, clear rebounds of tumorigenesis in the hIL-2 treated mice were evident (**Fig 6a**), indicating the potential benefits of the sustainable antitumor activity of DL_80_ for improving melanoma treatment.

**Figure 6.**
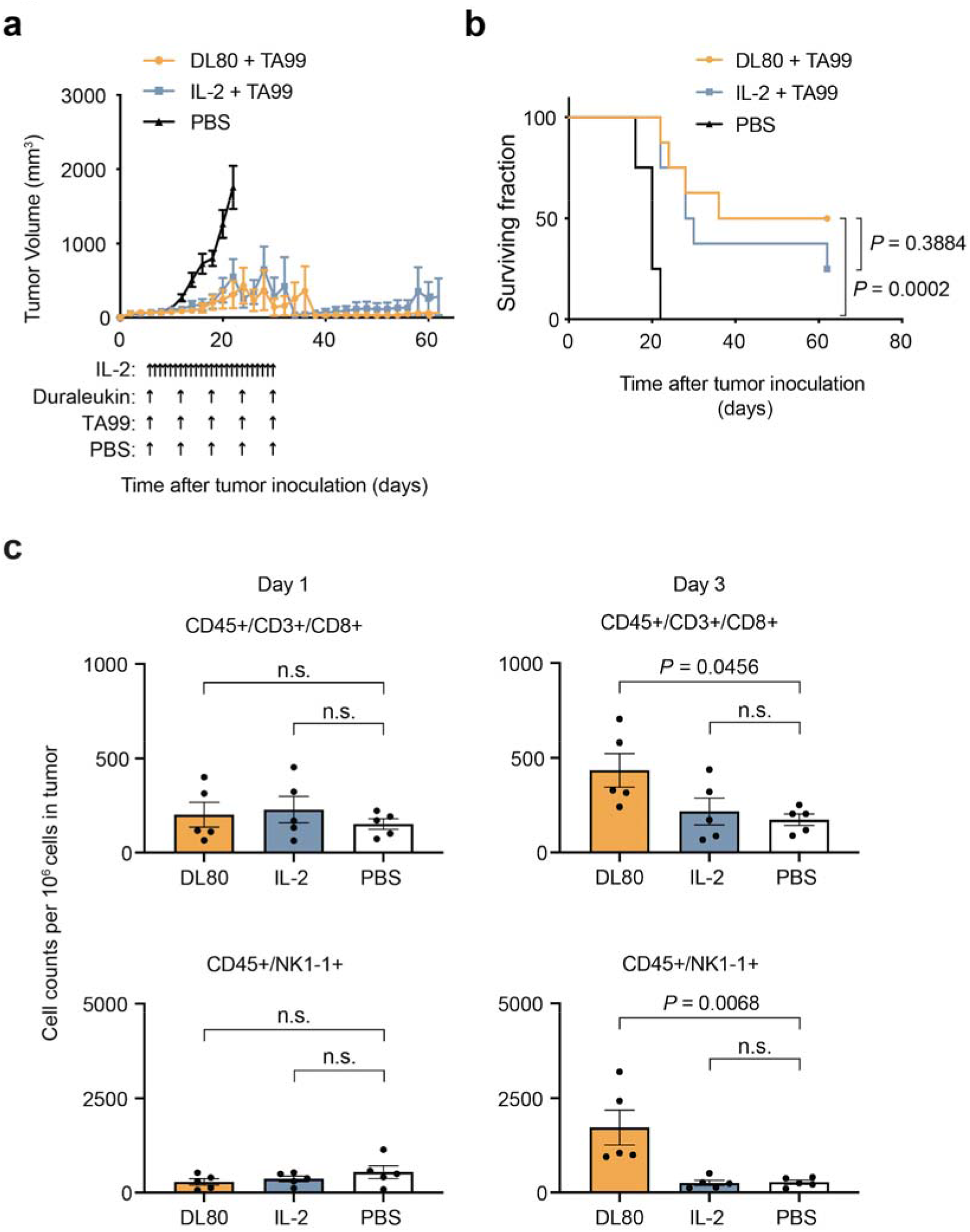
Evaluation of the *in vivo* efficacy of Duraleukin in a melanoma mouse model. a) The tumor growth curve. C57BL/6J mice bearing subcutaneous B16F10 tumors were treated with a combination of TA99 and Duraleukin or hIL-2 (n = 8/group) with different dosage intervals. PBS treatment was used for control. b) Animal survival curve after treatment. c) Flow cytometry analysis of tumor-infiltrating immune cells (n = 5/group). The y-axis is the respective immune cell count based on a million total cells from the isolated tumor.

To explore the mechanisms that contribute to Duraleukin’s superior antitumor activity, we isolated tumors at different stages and analyzed their immune cells by flow cytometry (**Fig 6c, Fig S10**). We observed significant and persistent increases of tumor-infiltrating immune cells, including CD8^+^ T cells and natural killer (NK) cells in the DL_80_-treated tumor tissues at day three post-treatment. At the same time, hIL-2 appeared to lose its activity for promoting T cell expansion. Our results indicate that Duraleukin exerts its long-lasting antitumor activity by promoting tumor-infiltrating immune cells.

## Discussion

The short serum half-lives limit the therapeutic and diagnostic efficacy of many small biotherapeutics, including cytokines, hormones, and Nbs. Here we develop a robust platform based on a large, diverse set of high-quality Nb_HSA_ to improve PK. We have employed and developed multidisciplinary approaches that span biochemistry, biophysics, structural proteomics, and integrative structural biology, longitudinal blood proteomics, and a quantitative proteomics-based PK assay that collectively enabled systematic characterizations of Nbs for advanced biomedical applications. We have positively evaluated the outstanding PK of Nb_HSA_ in a humanized HSA mouse model. To demonstrate the utility of this Nb platform, we applied several cross-species Nb_HSA_ to a key cancer-fighting cytokine (IL-2) and developed a specific Nb-based treatment (Duraleukin) for cancer. We demonstrated Duraleukin’s increased antitumor activity compared to IL-2 therapy alone in a melanoma mouse model.

The Nb platform that we have developed here likely represents a significant step towards tailored, optimal drug delivery. First, it permits minimal interference with the bioactivity of the small target drug compared with other approaches such as chemical modifications and direct fusion to HSA ^1,5^. The robustness of Nb_HSA_ is desirable for bioengineering. Second, it comprises a set of well-characterized Nb_HSA_ of excellent solubility, stability, and affinity. Their favorable physicochemical and structural properties are critical for drug development, administration, manufacturing, and storage ^12^. Rapid bioengineering and cost-effective production of Duraleukins, as demonstrated here, would be important for the future development of duraleukin technology towards clinical applications ^32^. Third, it offers a wide range of PK, biophysical and structural properties towards possible tailored drug development. Fourth, it does not elicit Fc-associated antibody-dependent cellular cytotoxicity (ADCC) that is likely toxic to the functional cells by which the Nb-fusion constructs target (such as IL-2R^+^ T cells). Combined with the high sequence similarity of Nbs to human IgGs, our approach may restrain the toxicity for clinical applications ^9,39^.

Our investigation leveraged a powerful combination of proteomic technologies for drug development. The integrative structural analysis revealed major HSA epitopes and provided a structural basis to understand Nb_HSA_ interactions with HSA and their PKs. Our analysis underlies the importance of high-affinity, acidic pH-dependent binding of Nb_HSA_ for half-life extensions. Our results indicate the presence of low intracellular concentration(s) of HSA-FcRn complex *in vivo* and the acidic microenvironment(s) of the FcRn receptor-mediated transcytosis ^3,4,40^. Moreover, we found that the PK of our most stable Nb_HSA_ is consistent with the half-life of HSA in the humanized mouse model. It is anticipated that Nb_HSA_ will have significantly improved PK in the non-human primate models and humans.

We developed Duraleukins by taking advantage of the cross-species Nb_HSA_ that help translate the preclinical data of animal models into clinical studies. Despite using a much smaller administration dose than hIL-2, Duraleukin_80_ outperformed it for melanoma treatment. The improved drug efficacy of Duraleukin is likely due to the robust cytokine activity and improved half-life. The retention of albumin in the tumor interstitium has been reported ^3,41,42^, raising a possibility that the superior efficacy of Duraleukin is related to its potentially favorable tumor localization as a cargo of albumin.

This proof-of-concept might be broadly applicable to other small antitumor cytokines such as hIL-15 ^43^, ultimately leading to enhanced antitumor activity, optimal dosing regimen, better drug safety, and improved patient compliance. It is conceivable that a combination of Duraleukin and an immune checkpoint blockade therapy like PD-1/L1 or CTLA-4 may further improve cancer treatment in a synergistic manner ^44,45^. It is also feasible to produce a trifunctional construct by adding another Nb that binds a prominent tumor surface marker (e.g., EGFR and HER2) for a more target-specific, and potentially more efficient, and safer therapy ^46^. Future studies will be needed to investigate the Nb_HSA_ and Duraleukin for preclinical and clinical applications.

## Supplemental Tables

**Table S1:** 71 Nb_HSA_ sequences, the pI, hydrophobicity, and the number of cysteine residues of Nb_HSA_ CDR3s, and the ELISA affinities of Nb_HSA_ towards different albumin species.

**Table S2**: SPR kinetic analysis, thermostability, and pharmacokinetic analysis of 20 different Nb_HSA_.

**Table S3**: Cross-link information and the cross-link models of the HSA-Nb_HSA_ complexes.

**Table S4**: Ion information of MS1-based and PRM quantification of Nb_HSA_.

**Figure S1.**
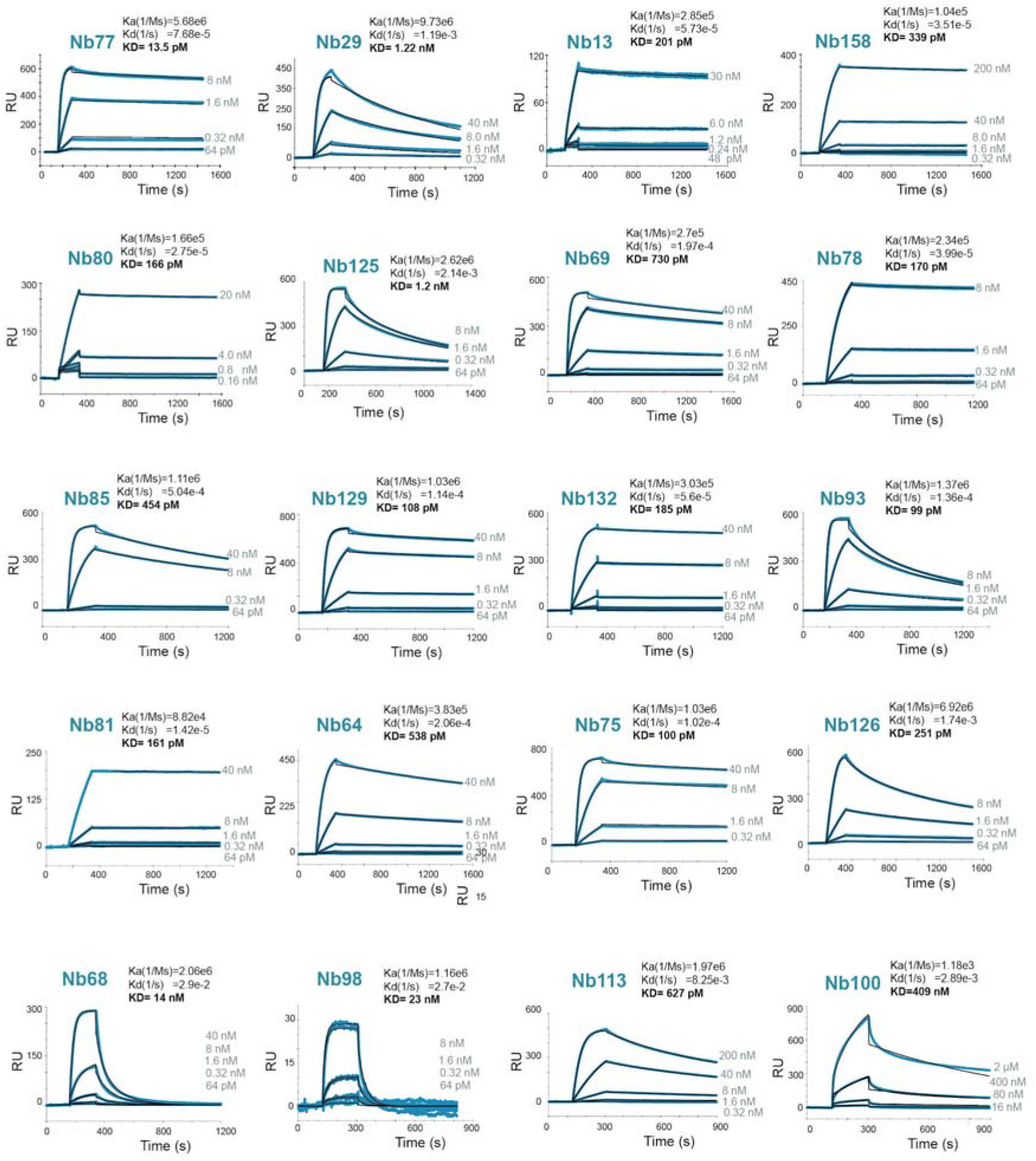
SPR affinity measurement of 20 Nb_HSA_. HSA binding kinetics of 20 Nb_HSA_ was measured by SPR (Biacore 3000). K_d_, dissociation rate constant; K_a_, association rate constant; K_D_, equilibrium dissociation constant.

**Figure S2.**
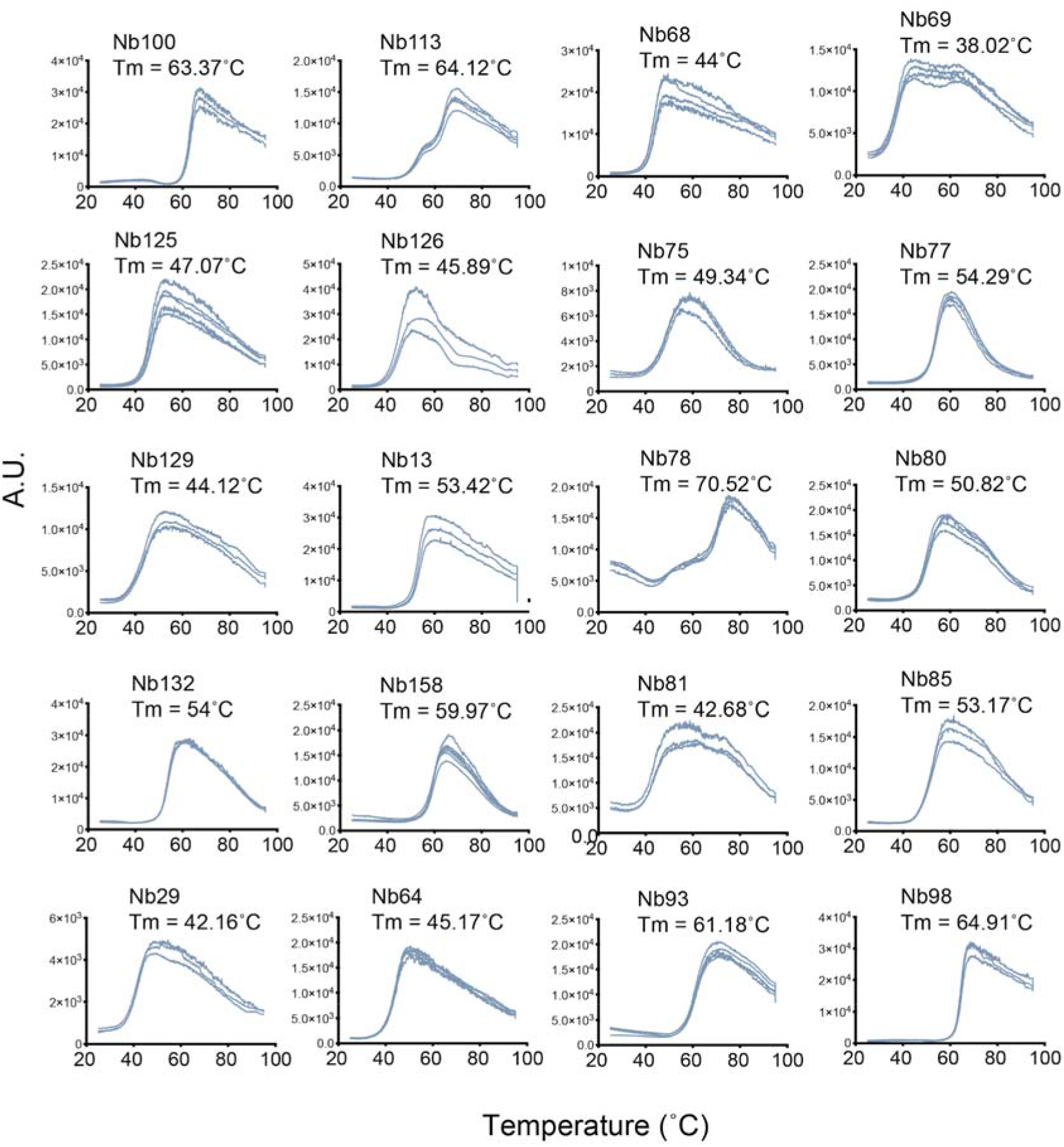
Thermostability measurements of 20 HSA-Nbs. The thermostability of 20 Nb_HSA_ was measured by differential scanning fluorimetry. Each measurement was repeated 3-6 times. T_M_: melting temperature. A.U.: arbitrary unit.

**Figure S3.**
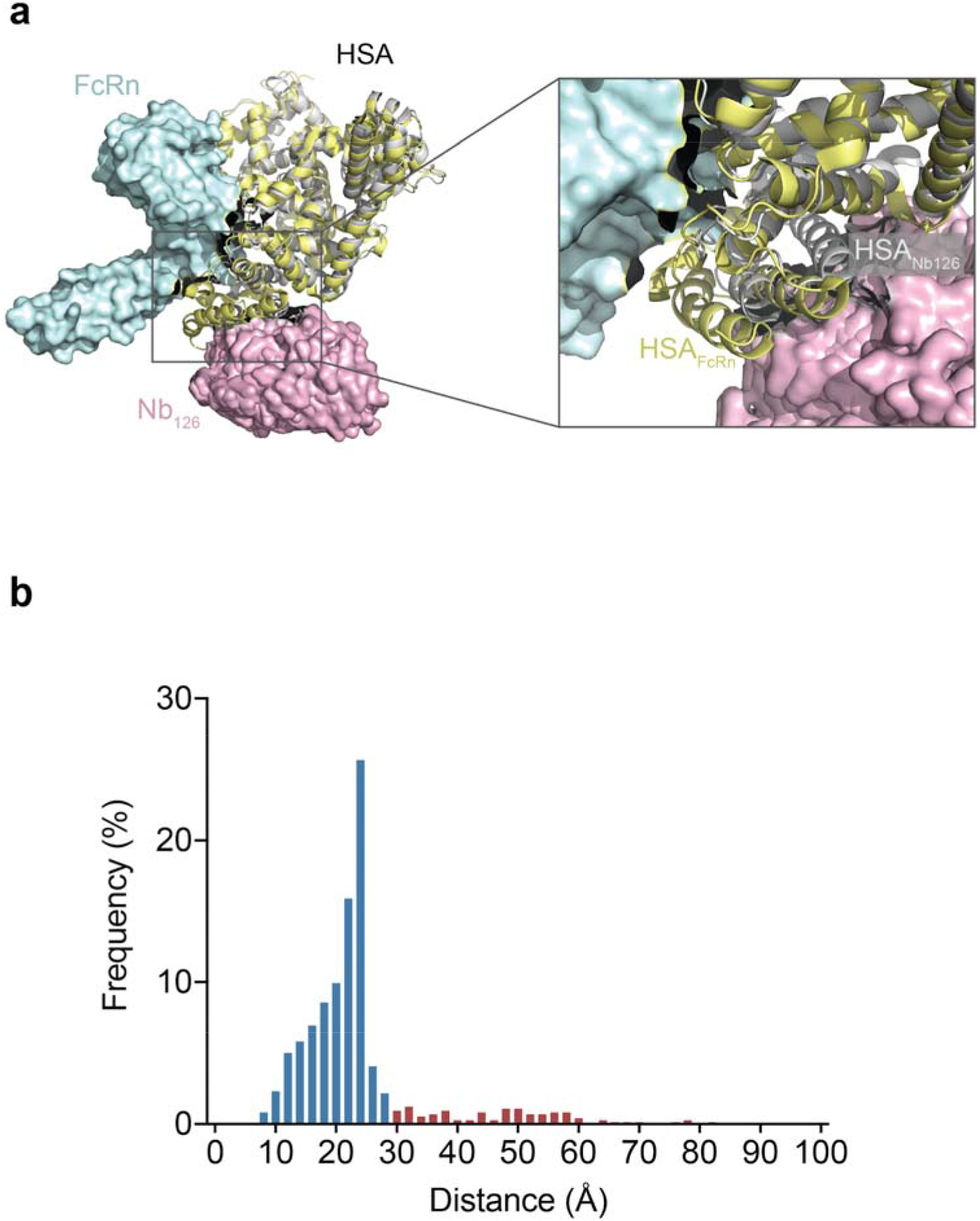
Overlay of FcRn binding HSA and Nb binding HSA structural models. a) Overlay of a crystallographic model of FcRn-HSA complex (PDB 4N0F) and the top 4 converged cross-link models of the Nb_126_-HSA complex. HSA is presented in cartoon style. Nbs and FcRn are presented as surfaces. Yellow: HSA bound with FcRn; blue: FcRn; grey: HSA bound with Nbs. b) The plot of the distances of the HSA-Nb intersubunit cross-links that are measured on the converged cross-link models.

**Figure S4.**
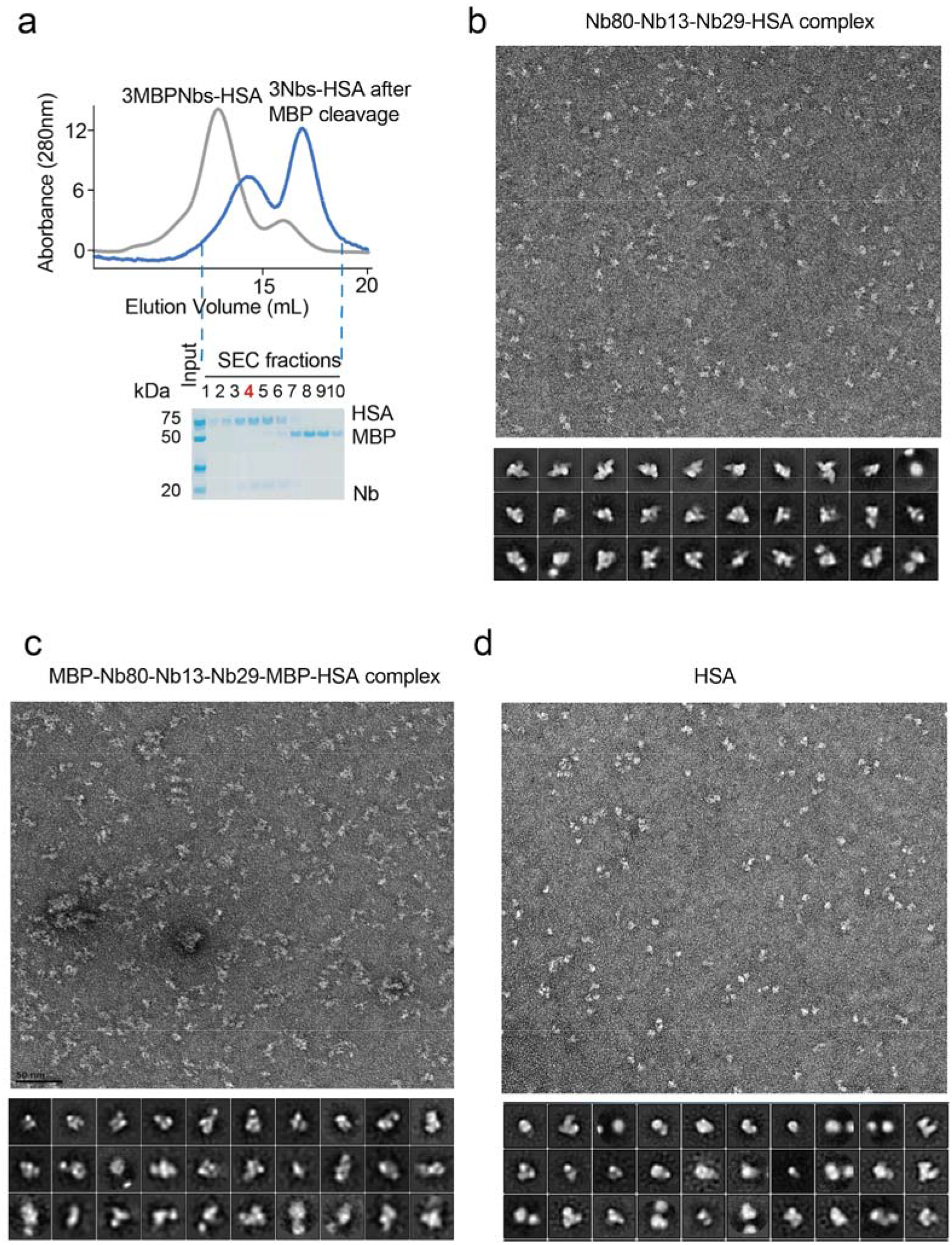
Purification and particle selection of tetrameric Nb-HSA complex. a) Purification of a homogenous HSA-Nb_80_-Nb_13_-Nb_29_ complex after removal of MBP tag by TEV protease cleavage. Size-exclusion chromatography and SDS-PAGE analysis confirmed the successful separation of the cleaved and uncleaved complexes. The homogenous SEC fraction four was used for negative stain EM analysis. b) Representative negative stain EM images of the purified tetrameric complex without the MBP tag. c) Representative negative stain EM images of the tetrameric complex in which each Nbs was fused with an MBP tag. d) Representative negative stain EM particles of HSA.

**Figure S5.**
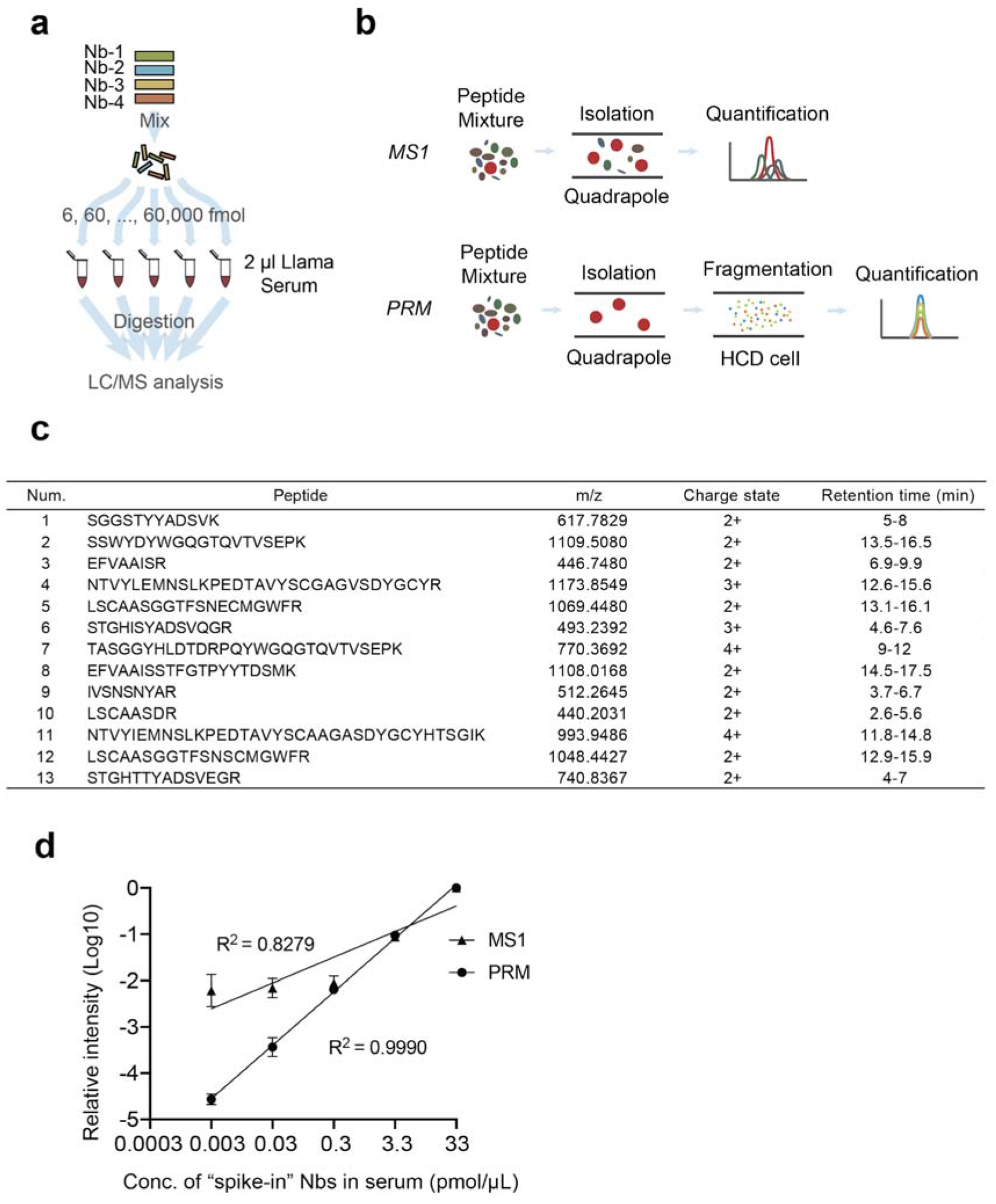
Development of an MS-based assay for multiplex Nb PK analysis. a) Schematic design of the method development. 4 different Nbs were first mixed in equal molarity. A ten-fold serial dilution of the Nbs from 60 pmole to 6 fmole was added to 2 μl llama serum. Sera proteins including the spike-in Nbs were proteolyzed. 1/1000 of the proteolyzed peptides of each sample was loaded onto the column and analyzed by LC/MS. b) Schematic comparison of MS1-based and MS2-based label-free quantification. c) Selected peptide information for MS1 quantification. d) Comparisons of the quantification linearity and sensitivity between MS1- and MS2-based label-free quantification approaches. Five orders of magnitude of quantification linearity (R^2^ = 0.999) was detected by PRM.

**Figure S6.**
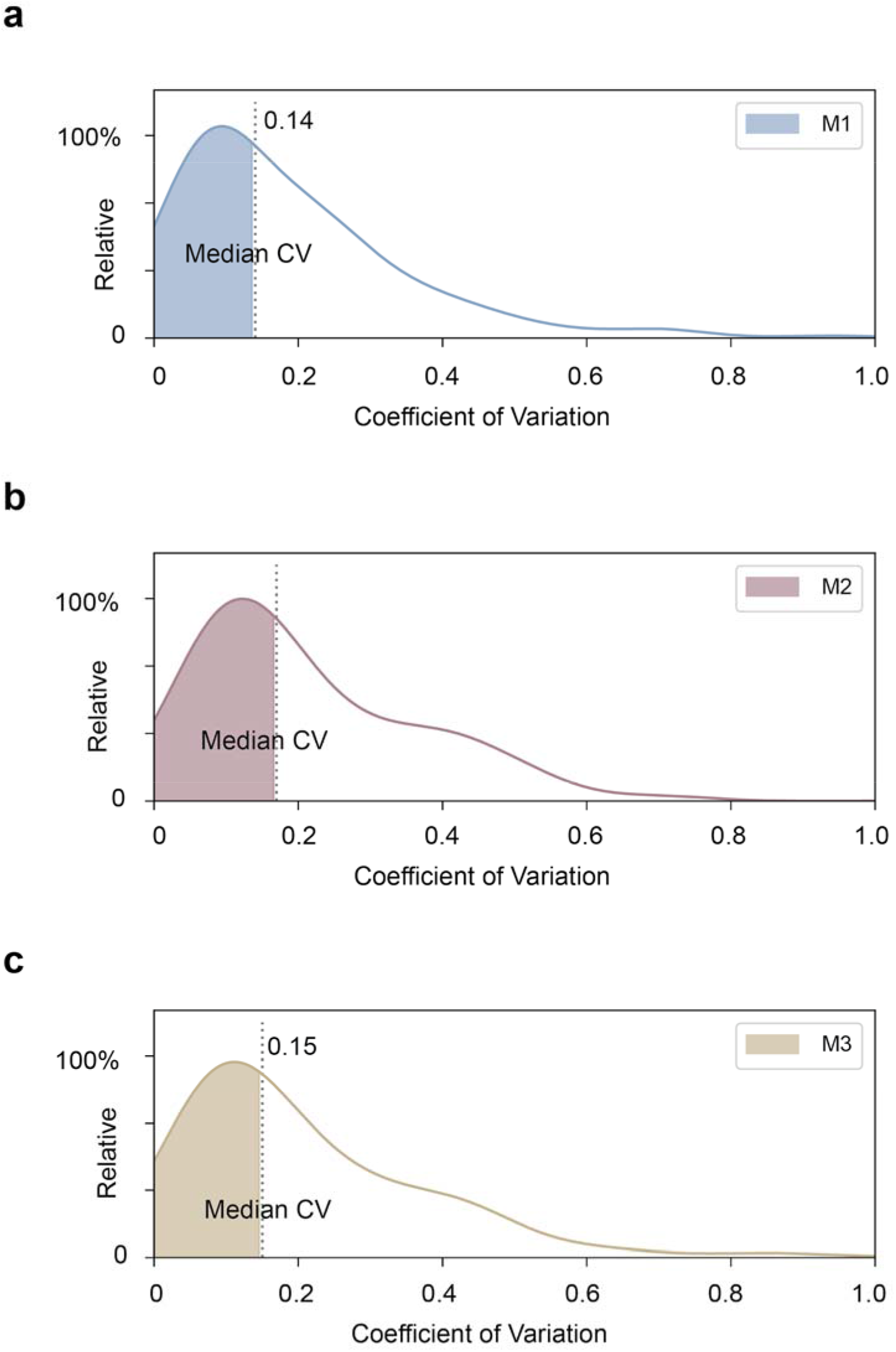
Coefficient of variation of *in vivo* Nb quantification. The coefficient of variations (CV) of Nb proteomic quantifications was plotted based on three technical replicates. M1- M3 indicates the measurements from three different animals.

**Figure S7.**
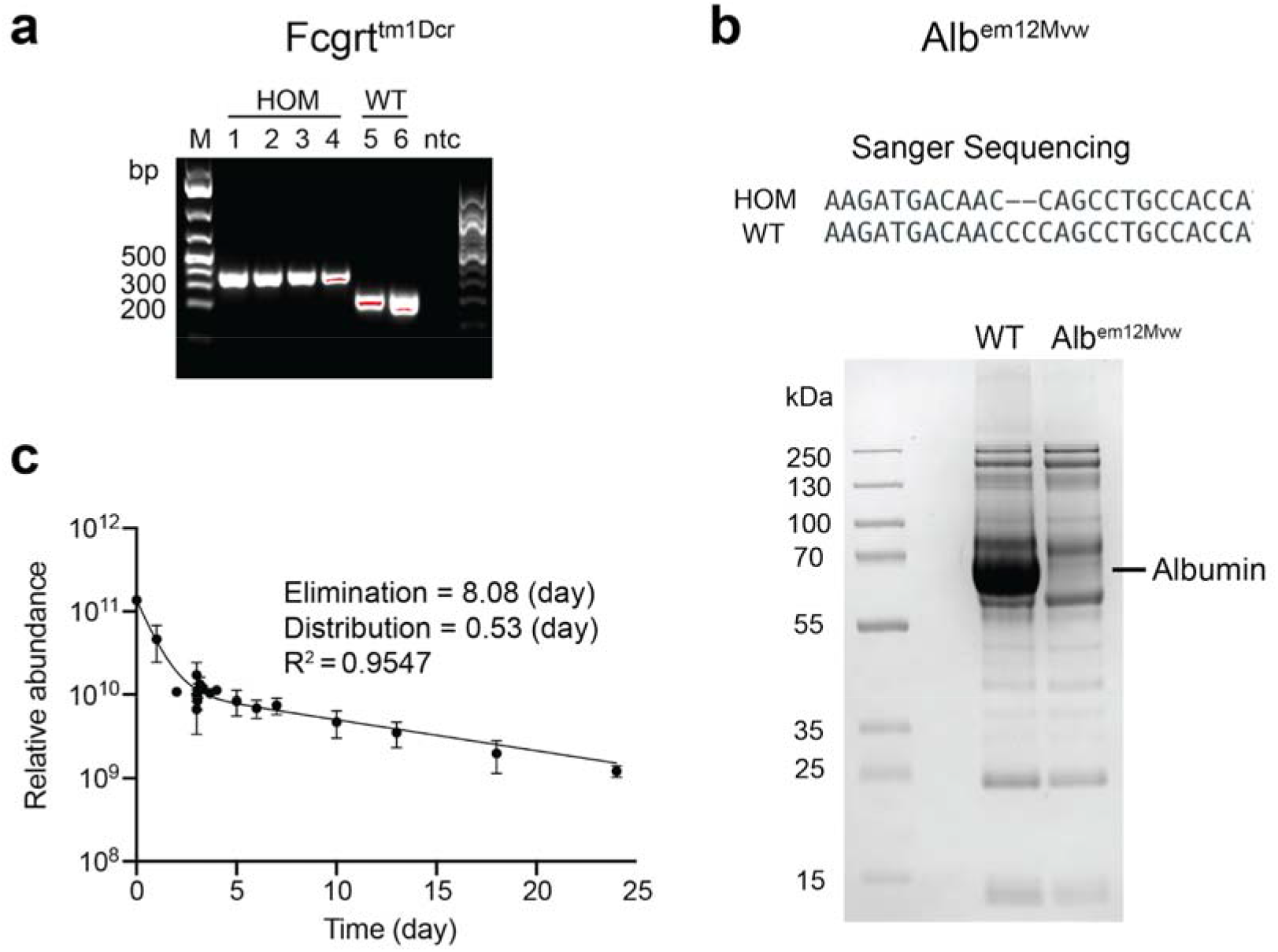
Verification of B6.Cg-Tg(FCGRT)32Dcr *Alb*^*em12Mvw*^ *Fcgrt*^*tm1Dcr*^/MvwJ mouse model. a) Verification of the *Fcgrt*^*tm1Dcr*^ homozygous strain (human FcRn knock-in) by DNA electrophoresis. HOM: homozygous mouse. b) Verification of the *Alb*^*em12Mvw*^ strain by Sanger sequencing (upper panel) and SDS-PAGE (phenotypic) analysis (lower panel). WT: C57BL/6J strain; Alb^em12Mvw^: albumin knock-out strain. c) Pharmacokinetics of HSA in the mouse model measured by label-free LC/MS.

**Figure S8.**
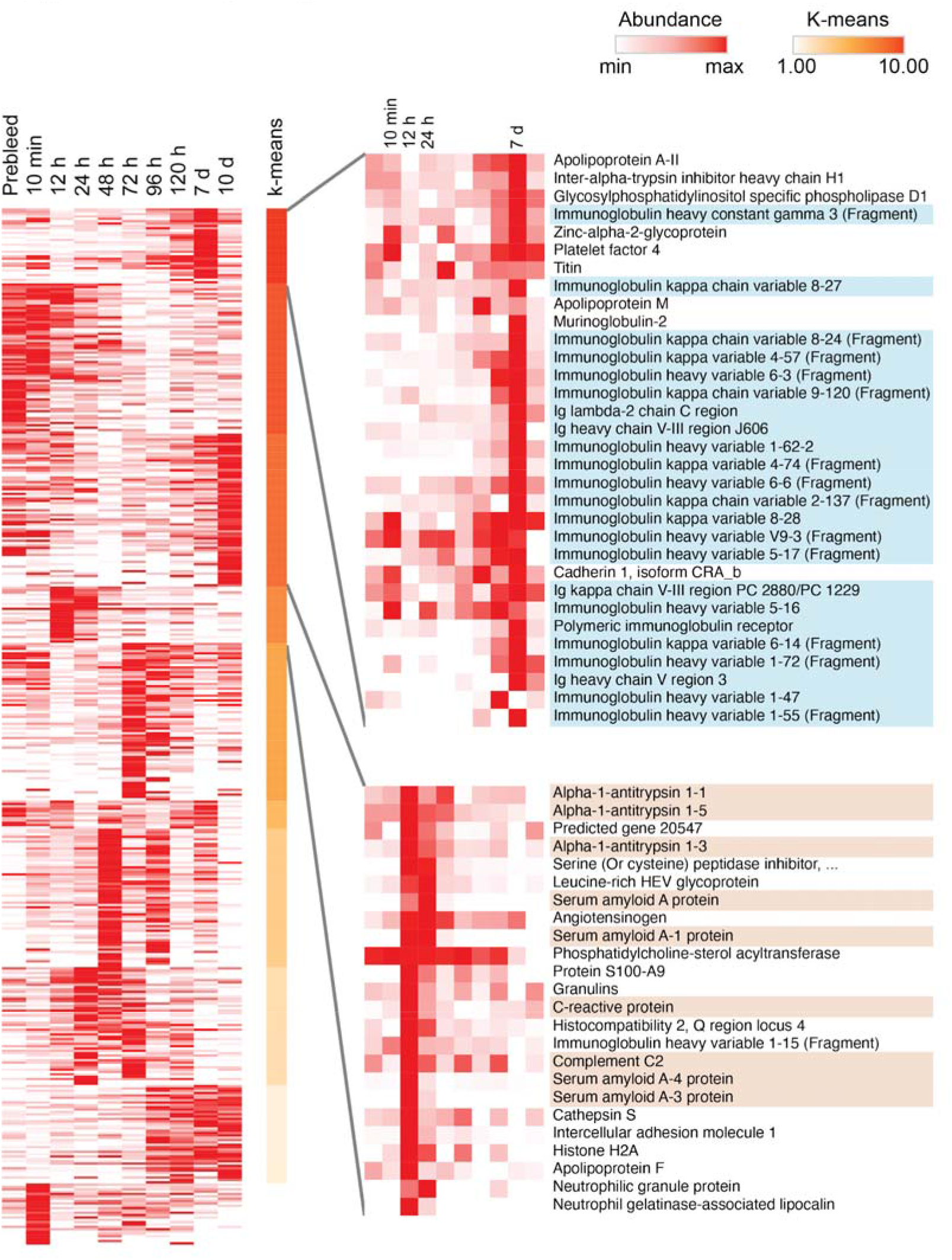
Serum protein quantification upon HSA administration in the humanized mouse model. Mouse blood was collected at different time points. The resulting serum proteomes were quantified by label-free LC/MS. The relative abundances of individual proteins among ten different time points were quantified and clustered based on the K-means method. Serum protein groups peaked at 12-24 hours (e.g., acute-phase inflammatory proteins) or at day 7 (immunoglobulins) were zoomed in.

**Figure S9.**
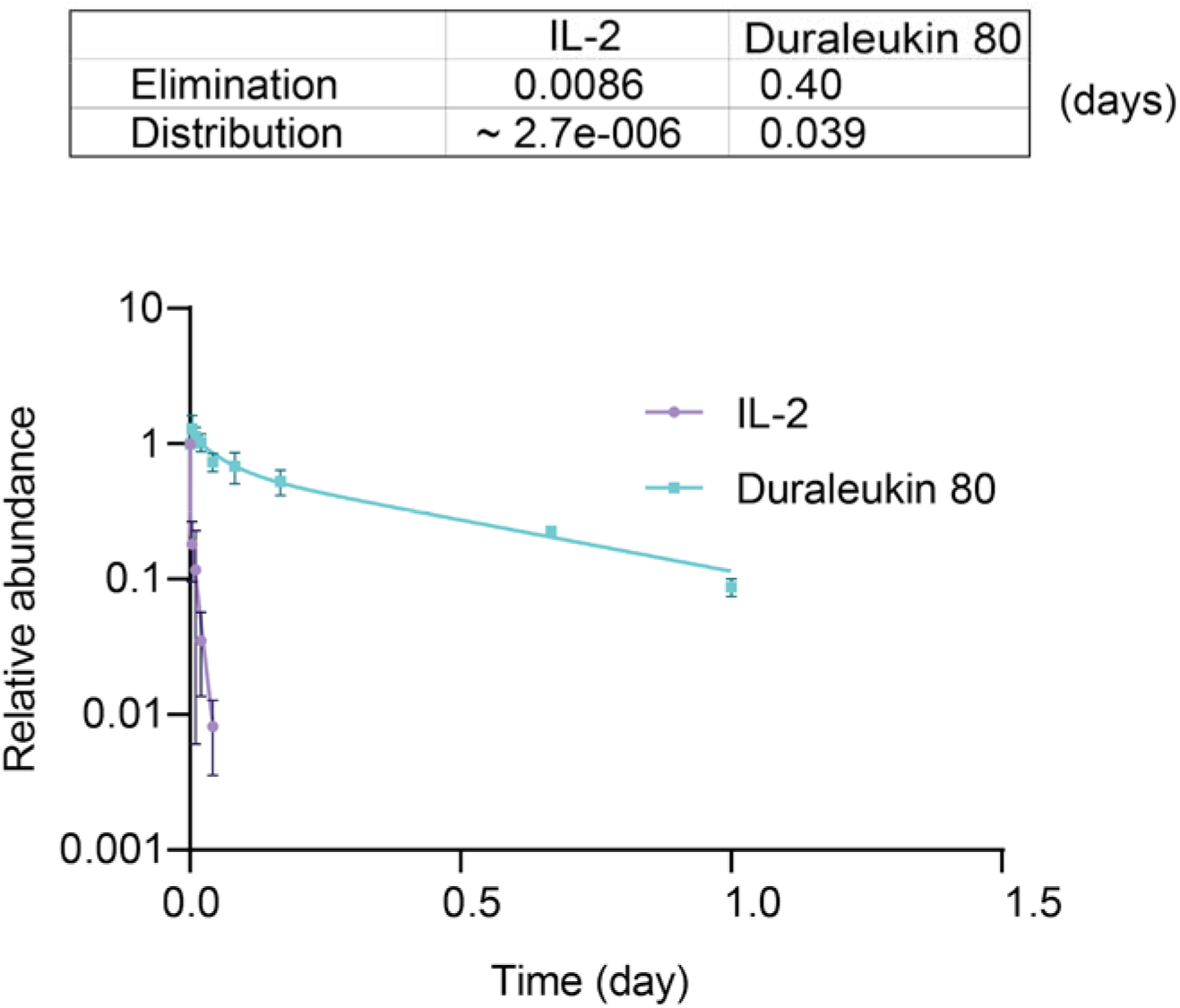
Pharmacokinetics (day) of DL_80_ in C57BL/6J mouse. C57BL/6J mice were treated with DL_80_ or hIL-2 (equimolar). Blood was sampled at different times (up to 1 day) and was collected for ELISA measurements using an anti-hIL-2 monoclonal antibody (BG5). The distribution and elimination rates (day) were calculated based on the ELISA PK curves by fitting into a two-phase decay model (**Methods**).

**Figure S10.**
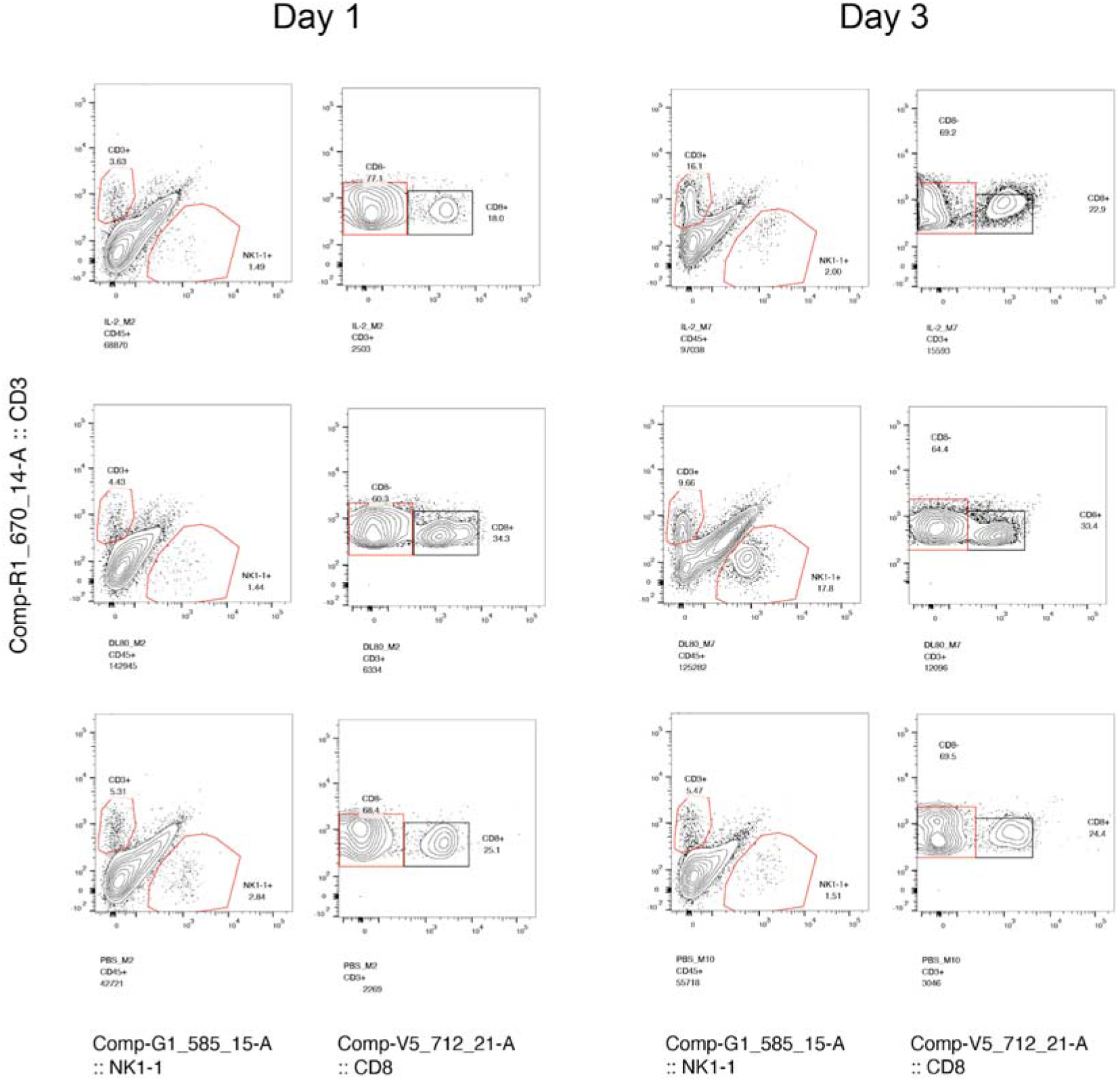
Representative flow cytometry analyses of the CD8^+^ T cells and NK cells isolated from the mouse melanoma post-treatment by either Duraleukin, IL-2, or PBS. Melanoma tumor tissues were isolated from the animals at day one and day three post-treatment by either Duraleukin (DL), IL-2, or PBS control. A mixture of tumor cells and immune cells were isolated, and the samples were stained by antibodies that recognize specific immune cell markers before analysis by flow cytometry. IL-2 treated samples: upper panel; DL-treated samples: middle panel; PBS-treated samples: lower panel. Left: day 1 post-treatment; Right: day 3 post-treatment. n = 5/group.

## Methods

### Nb DNA synthesis and cloning

Nb genes were codon-optimized for expression in *Escherichia coli* and the nucleotides were *in vitro* synthesized (Synbio). The Nb genes were cloned into a pET-21b (+) vector at BamHI and XhoI restriction sites, or a pMAL-c5X vector at BamHI and EcoRI restriction sites.

### Nb Purification

Nb DNA constructs were transformed into BL21(DE3) cells and plated on Agar with 50 μg/ml ampicillin at 37 °C overnight. A single bacterial colony was picked and cultured in LB broth to reach O.D. at ~ 450 nm. 0.1 or 0.5 mM IPTG was added to the *E. coli* cell culture for MBP fusion Nbs or his-tagged Nbs induction at 16°C overnight. Cells were then harvested, briefly sonicated, and lysed on ice with a lysis buffer (1xPBS, 150 mM NaCl, 0.2% TX-100 with protease inhibitor). After lysis, soluble protein extract was collected at 15,000xg for 10 mins, and recombinant his-tagged Nbs were purified by His6-Cobalt resin (Thermo) and eluted by imidazole; while MBP-fusion Nbs were purified by amylose resin (NEB) and eluted by maltose. The eluted Nbs were subsequently dialyzed in the dialysis buffer (e.g., 1x DPBS, pH 7.4). A HiLoad 16/600 Superdex 200 pg column (GE Healthcare) on an ÄKTA FPLC protein purification system (GE Healthcare) was used to further purify the Nbs and ensure their monomeric states. The purified Nbs were analyzed by SDS-PAGE. For the animal experiments, excessive endotoxin in the Nbs was removed (below 0.1 EU/dose) by the oxinEraser Endotoxin Removal column (Genscript) and measured by ToxinSensor Chromogenic LAL Endotoxin Assay Kit (Genscript). The purified Nbs were further sterilized by passing a 0.2 μm filter (Millex) and were stored at −80°C before use.

### ELISA (Enzyme-linked immunosorbent assay)

Indirect ELISA was carried out to measure the relative affinities of Nbs. Albumin was coated onto a 96-well ELISA plate (R&D system) at 1-10 ng/well in coating buffer (15 mM sodium carbonate, 35 mM sodium bicarbonate, pH 9.6) overnight at 4°C and was blocked with a blocking buffer (DPBS, 0.05% Tween 20, 5% milk) at room temperature for 2 hrs. Nb_HSA_ were diluted in the blocking buffer and incubated with albumin for 2 hrs. HRP-conjugated secondary antibodies against His-tag (Genscript) were diluted 1:5,000-10,000 and incubated with the well for 1 hr at room temperature. After PBST (DPBS, 0.05% Tween 20) washes, the samples were further incubated under dark with freshly prepared w3,3′,5,5′-Tetramethylbenzidine (TMB) substrate for 10 mins to develop the signals. After the STOP solution (R&D system), the plates were read at multiple wavelengths (the optical density at 550 nm wavelength subtracted from the density at 450 nm) on a plate reader (Multiskan GO, Thermo Fisher). The relative affinity of each Nb was determined by the average of O.D. readouts. For human and monkey albumin, relative affinity of each Nb was tested at Nb concentration of 1 μM, 100 nM, 10 nM, 1 nM, and 100 pM. An average ELISA O.D. of >0.1 was used as the cutoff value to assign albumin binding Nbs. For mouse albumin, relative affinity was tested at one μM and a cutoff O.D. of >1 was used.

### *In vitro* pull-down assay (Immunoprecipitation)

Nb or serum albumin was coupled to CNBr-activated sepharose resin (GE Healthcare). For the i*n vitro* pull-down assay, different concentrations of Nbs were incubated with the human serum albumin coupled resin. Samples were incubated for 15-30 min at 4°C with gentle agitation. The resin was collected and washed three times with a washing buffer and was boiled in the LDS sample loading buffer (Thermo) before SDS PAGE analysis.

### Nb Affinity measurement by surface plasmon resonance (SPR)

Surface plasmon resonance (SPR, Biacore 3000 system, GE Healthcare) was used to measure Nb affinities. Briefly, human serum albumin was immobilized to the flow channels of an activated CM5 sensor-chip. Protein analytes were diluted to 10 μg/ml in 10 mM sodium acetate, pH 4.5, and injected into the SPR system at 5 μl/min for 420 s. The surface was then blocked by 1 M ethanolamine-HCl (pH 8.5). For each Nb analyte, a series of dilution (spanning ~ 1,000-fold concentration range) was injected in duplicate, with HBS-EP+ running buffer (GE-Healthcare) at a flow rate of 20- 30 μl/min for 120- 180 s, followed by a dissociation time of 10 – 20 mins based on dissociation rate. Between each injection, the sensor chip surface was regenerated twice with a low pH buffer containing ten mM glycine-HCl (pH 1.5- 2.5) at a flow rate of 40-50 μl/min for 30 s - 1 min. Binding sensorgrams for each Nb were processed and analyzed using BIAevaluation by fitting with the 1:1 Langmuir model.

### Clustering and phylogenetic tree analysis of Nb_HSA_

A phylogenetic tree was generated by Clustal Omega ^47^ with the input of unique Nb_HSA_ CDR3 sequences and the adjacent framework sequences (i.e., YYCAA to the N-terminus and WGQG to the C-terminus of CDR3s) to help alignments. The data was plotted by ITol (Interactive Tree of Life) ^48^. Isoelectric points and hydrophobicities of the CDR3s were calculated using the BioPython library. The sequence logo was plotted using WebLogo ^49^.

### Thermostability analysis of Nb_HSA_

The thermal stability of Nb_HSA_ was measured by differential scanning fluorimetry (DSF). To prepare DSF samples, Nb_HSA_ were mixed with SYPRO orange dye (Invitrogen) in PBS to reach a final concentration of 2.5-15 μM. The samples were analyzed by a 7900HT Fast Real-Time PCR System (Applied Biosystems) in triplicate. The temperature was programmed to increase from 25°C to 95°C with a ramp rate of 1°C/min to generate the melting curves. The melting point was calculated by first derivatives method ^50^.

### Chemical cross-linking and mass spectrometry (CXMS)

Nb was incubated with albumin in PBS and 1 mM dithiothreitol (DTT) to allow the formation of the complex. HSA-Nb_HSA_ complexes were crosslinked with 1 mM disuccinimidyl suberate (DSS, ThermoFisher Scientific) for 23 min at 25°C with gentle agitation. The crosslinking reaction was then quenched with 50 mM ammonium bicarbonate (ABC) for 10 min at room temperature. After protein reduction and alkylation, the cross-linked samples were separated by a 4–12% SDS-PAGE gel (NuPAGE, Thermo Fisher). The regions corresponding to the cross-linked species (~130 kDa) were cut and in-gel digested with trypsin and Lys-C as previously described ^16,19,51^. After proteolysis, the peptide mixtures were desalted and analyzed with a nano-LC 1200 (Thermo Fisher) coupled to a Q Exactive™ HF-X Hybrid Quadrupole-Orbitrap™ mass spectrometer (Thermo Fisher). The cross-linked peptides were loaded onto a picochip column (C18, 3□μm particle size, 300□Å pore size, 50□μm□×□10.5□cm; New Objective) and eluted using a 60□min LC gradient : 5% B–8% B, 0 – 5 min; 8% B – 32% B, 5 – 45 min; 32% B–100% B, 45 – 49 min; 100% B, 49 −54 min; 100% B - 5 % B, 54 min - 54 min 10 sec; 5% B, 54 min 10 sec - 60 min 10 sec; mobile phase A consisted of 0.1% formic acid (FA), and mobile phase B consisted of 0.1% FA in 80% acetonitrile. The QE HF-X instrument was operated in the data-dependent mode, where the top 8 most abundant ions (mass range 380–2,000, charge state□3 - 7) were fragmented by high-energy collisional dissociation (normalized collision energy 27). The target resolution was 120,000 for MS and 15,000 for MS/MS analyses. The quadrupole isolation window was 1.8□Th, and the maximum injection time for MS/MS was set at 120□ms. After the MS analysis, the data was searched by pLink for the identification of cross-linked peptides. The mass accuracy was specified as 10 and 20□p.p.m. for MS and MS/MS, respectively. Other search parameters included cysteine carbamidomethylation as a fixed modification and methionine oxidation as a variable modification. A maximum of three trypsin missed-cleavage sites was allowed. The initial search results were obtained using the default 5% false discovery rate, estimated using a target-decoy search strategy. The crosslink spectra were then manually checked to remove potential false-positive identifications essentially as previously described ^16,19,51,52^.

### Integrative structural modeling of HSA-Nb complexes

Structural models for Nbs were obtained using a multi-template comparative modeling protocol of MODELLER^53^. Next, we refined the CDR3 loop and selected the top 5 scoring loop conformations for the downstream docking. Each Nb model was then docked to the HSA structure (PDB 4g03) by an antibody-antigen docking protocol of PatchDock software that focuses the search to the CDRs^54^ and optimizes CXMS-based distance restraints satisfaction^55^. A restraint was considered satisfied if the Ca-Ca distance between the cross-linked residues was within 30Å for DSS cross-linkers ^16,56^. The models were then re-scored by a statistical potential SOAP ^57^. The antigen interface residues (distance <6Å from Nb atoms) among the 10 best scoring models according to the SOAP score, were used to determine the epitopes. Convergence was measured as the average RMSD among the 10 top-scoring models. Once the epitopes were defined, we clustered the Nbs based on the epitope similarity using hierarchical clustering. The clusters reveal the most immunogenic surface patches on the antigens.

### Size exclusion chromatography (SEC) and negative stain electron microscopy

HSA and MBP-Nbs were mixed in a HSA:Nb ratio of 1:1.2 and incubated at room temperature for 2 hours in PBS and 2 mM DTT. The complexes were analyzed by SEC (Superdex200, GE LifeSciences). HSA and the tetrameric HSA-Nb_HSA_ complex were visualized by transmission electron microscopy (TEM) under negative staining. For TEM analysis, the complexes were digested overnight at 4 °C with TEV protease to remove MBP tag. Protein was diluted to a final concentration of 30 μg/ml (HSA in PBS; HSA-Nb complexes in Tris 20 mM, 200mM NaCl, 3% Glycerol) and deposited on carbon-coated CF400-CU grids (EMS) which were freshly glow discharged. After 30 seconds of incubation, the excessive protein was removed, and the grid was stained with two drops of uranyl acetate 2 % w/v. Electron micrographs were recorded in an FEI TECNAI T12 operating at 120 kV with a 2k x 2k Gatan UltraScan 1000. Raw images were converted by IMOD 4.8 ^58^and particles were selected using EMAN2 ^59^. Particle processing and 3D reconstruction were done by Relion 3.0 ^60^ with several rounds of auto-refine iterations. The final 3D structures were obtained with 4,000 particles for HSA and 21,400 particles for the tetrameric complex. Crystallographic structural models (PDB: 1AO6 and 5VNW) were used for the initial fitting by Chimera ^61^.

### Site-directed mutagenesis

An HSA expression plasmid was obtained from Addgene (ALB-bio-His, Plasmid #52176). E400R and K383D point mutations were introduced to the HSA sequence by the Q5 site-directed mutagenesis kit. After sequence verification by Sanger Sequencing, plasmids bearing wild type HSA and the mutants were transfected to HeLa cells using Lipofectamine 3000 transfection kit (Invitrogen) and Opti-MEM (Gibco) according to the manufacturer’s protocol. The cells were cultured overnight before a change of medium to DMEM without FBS supplements to remove BSA. After a 48 h culture at 37°C, 5% CO_2_, the media expressing HSA were collected and stored at −20°C. The media were analyzed by SDS-PAGE and Western Blotting to confirm protein expression.

### MS quantification of Nb pharmacokinetics

600 μg/g of HSA (w:w) was administered in B6.Cg-Tg(FCGRT)32Dcr *Alb*^*em12Mvw*^ *Fcgrt*^*tm1Dcr*^/MvwJ mice (JAX, n = 3) by *intravenous (i.v.)* injection three days before the injection of Nbs. A mixture of Nbs including 20 Nb_HSA_ and 2 control Nbs in equal molarity were then injected at 30 μg/g (w:w, day 0). After injections, the whole blood samples were collected from the tail veins of the mice. The blood sampling time for HSA was pre-dose (0 min), 10 min, 24 h, 48 h. The sampling time for Nb mixture was pre-dose (0 min), 5, 15, 30 min, 1, 2, 4, 8, 16, 24 hrs, 2, 3, 4, 7, 10, 15, 21 days post-dose. Whole blood samples were allowed to clot at room temperature for 30 min and were centrifuged at 4°C, 20,000 × g for 3 min. Serum proteins from the supernatants were collected for the downstream proteomic analysis.

Serum samples were reduced in 8M urea digestion buffer (with 50 mM Ammonium bicarbonate, 5 mM TCEP, and DTT) at 57°C for 1 hr, and alkylated in the dark with 30 mM Iodoacetamide for 30 mins at room temperature. The samples were in-solution digested with 1:100 (w/w) trypsin and Lys-C at 37°C overnight, with incubation with another bolus of trypsin 4 hours. After proteolysis, the peptide mixtures were desalted and analyzed with a nano easyLC 1200 device coupled with a Q Exactive™ HF-X Hybrid Quadrupole Orbitrap™ mass spectrometer (Thermo Fisher). Briefly, Nb peptides were loaded onto a Picochip column (C18, 1.9 μm particle size, 120□Å pore size, 75□μm□×□25□cm; New Objective) and eluted using a 45-min liquid chromatography gradient (5% B–8% B, 0–3 min; 8% B–42% B, 3–35 min; 42% B–100% B, 35 – 40 min; 100% B, 40 - 45 min; mobile phase A consisted of 0.1% formic acid (FA), and mobile phase B consisted of 0.1% FA in 80% acetonitrile (ACN)). The flow rate was ~300 nl/min. The QE HF-X instrument was operated in the data-dependent mode, where the top 6 most abundant ions (mass range 300 – 2,000, charge state□2 - 8) were fragmented by high-energy collisional dissociation (HCD). The target resolution was 120,000 for MS and 60,000 for MS2 analyses. The quadrupole isolation window was 1.8□Th, and the maximum injection time for MS/MS was 120□ms.

Serum proteins including Nbs were first identified by MaxQuant version 1.6.1.0 ^62^. Up to 4 unique and specific tryptic peptides with the preference of CDR3-containing peptides with high intensities were selected from each Nb.

For the MS1-based quantification, the area under the curve (AUC) of the selected peptides were calculated from the raw data by using Xcalibur Qual Browser. For each Nb, AUCs of the selected peptides were averaged to represent the abundance of the Nbs. Selected peptides were validated by MS/MS.

For the MS2-based quantification, an inclusion list of the Nb peptides to be monitored was generated. An LC RT window of −2~+3 min (for a 90-min LC gradient) was used to ensure that all the targeted peptides were included. The instrument was operated in a data-independent model where the targeted peptides were specifically selected for fragmentation (HCD normalized energy 27). Up to 5 abundant fragment ions from each peptide were chosen to increase the specificity of our quantification. AUCs of the fragment ions were calculated using Xcalibur Qual Browser. The relative abundance of Nbs was presented as the median AUC intensities of their respective MS2 fragment ions. It was further normalized based on the total MS1 ion current (TIC) of each LC run. An in-house script was developed to enable automatic quantification described above. The samples were analyzed in triplicates. The abundance of serum Nbs in pharmacokinetics analysis was determined by averaging technical and biological replicates; the resulting data was used to fit into a two-phase decay model by Prism GraphPad.

### Serum proteomic quantification

1 mg HSA was administered to a B6.Cg-Tg(FCGRT)32Dcr *Alb*^*em12Mvw*^ *Fcgrt*^*tm1Dcr*^/MvwJ mouse. Blood samples were collected from the tail vein at the following time: pre-dose, 10 min, 24, 48, 72, 96 h, 7, 10 days. Serum sample preparation and in-solution digestion of serum proteins were processed, as previously mentioned. Peptide samples were analyzed with a nano easy□C 1000 device coupled with a Q Exactive™ HF-X Hybrid Quadrupole Orbitrap™ mass spectrometer (Thermo Fisher). Briefly, Nb peptides were loaded onto an analytic column (1.6 μm particle size, 100□Å pore size, 75□μm□×□25□cm; IonOpticks) and eluted using a 90-min liquid chromatography gradient. A QE HF-X instrument was used for analysis. The quadrupole isolation window was 1.6□Th, and the maximum injection time for MS/MS was set at 100□ms. Serum protein levels were quantified using MaxQuant version 1.6.1.0. The resulting raw data were searched against a mouse Uniprot FASTA protein database (December 2018) with fixed modification of carbamidomethylation and variable modifications of methionine oxidation and N-terminal acetylation. Trypsin was set as a digestion enzyme, and a maximal missed cleavage of 2 was allowed. Protein’s false discovery rate was set to be 0.01. Minimal peptide length was 7, and maximal peptide mass was 4600 Da. Minimal ratio count of label-free quantification was set to be 2. Serum protein abundance at each time point is derived by averaging ion intensities of 3 technical replicates and normalized based on baseline abundance. Serum proteins were then clustered using the K-means clustering algorithm (K=10), and heatmap was generated by software Morpheus (https://software.broadinstitute.org/morpheus).

### Purification of Duraleukins

hIL-2 (C125A) sequence was fused to C-termini of Nb sequences at the DNA level with a flexible linker (GGGGS)_2_ (Gly-Gly-Gly-Gly-Ser-Gly-Gly-Gly-Gly-Ser). The resulting DNA sequences were synthesized and cloned into pET-21b(+) (Synbio). Plasmids were transformed into BL21(DE3) *E. coli* cells to produce Duraleukins. After IPTG induction, Duraleukins were purified from the cell pallets. Cell pellets containing the inclusion bodies of Duraleukins were washed with a solubilization buffer A (0.1 M Tris, 2% sodium deoxycholate (SDC), 5 mM EDTA, pH 8) and sonicated on ice. Cell lysates were spun down at 12,000 × g for 20 min 4°C. Supernatants were discarded, and the pellets were saved for analysis. Cell pellets were washed with Milli-Q water and spun down at 12,000 × g for 20 min at 4°C. Solubilization buffer B (0.1 M Tris, 6 M guanidine hydrochloride (GuHCl), pH 8) was added to the pallets and incubated for 1 h incubation at room temperature with gentle agitation. After centrifugation 12,000 × g for 20 min at 4°C, the supernatants that contain soluble forms of Duraleukins were collected, and subsequently diluted with a Duraleukin refolding buffer (0.1 M Tris, pH 8, the final concentration of 10 mM reduced and 1 mM oxidized glutathione) to a GuHCl concentration of 2 M and a protein concentration of ~0.3 mg/mL. The solutions were then incubated at room temperature for 16 h with gently rotating to allow protein refolding. The insoluble fraction was removed by centrifugation at 16,000 × g for 20 min at 4°C, and refolded Duraleukins were obtained in the supernatants. IMAC and FPLC purification, detoxification, and sterilization were performed as previously described.

### In vitro T cell proliferation assay

CTLL-2 cells (ATCC) were cultured, according to ATCC’s protocol. The complete culture medium was formulated as 10% FBS, an additional 2 mM L-glutamine, and 1 mM sodium pyruvate in ATCC-formulated RPMI-1640 containing 10% T-STIM with Con A (Corning). For CTLL-2 proliferation assay, cells were grown to a density of 1 × 10^5^ cells/mL. Duraleukins or hIL-2 (PeproTech) were added into the wells of a flat-bottom 96-well tissue culture microplate. The final concentration of hIL-2 and Duraleukins was adjusted to 0.01~25 ng/mL and 0.025~50 ng/mL by 2-fold serial dilution. Every concentration group was tested in duplicate. 10^4^ cells were seeded to each well of the plate, and the total volume in each well was 100 μL. The cells were then incubated at 37°C, 5% CO_2_ for 56 h. 10 μL of WST-1 reagent (Roche) was added to each well and the cells were incubated for another four h. The plate was shaken at 1,000 rpm for 1 min before analysis. A spectrometer read absorbance at 450 nm. 690 nm was used as a reference wavelength. A dose-response curve was fit via nonlinear regression using GraphPad Prism 7.

### Generation and treatment of melanoma mouse model

B16F10 (ATCC) was cultured, according to ATCC’s protocol. The complete medium is formulated as 10% FBS in ATCC-formulated DMEM. C57BL/6J mice were purchased from The Jackson Laboratory. All mice were aged eight weeks before tumor inoculation. B16F10 melanoma cells were grown into the logarithmic growth phase (≤ 50% confluent), harvested, and adjusted to a concentration of 1 × 107 cells/ml in ice-cold HBSS. Mice were anesthetized with isoflurane and shaved in the middle of the back with a razor. 100 μl (10^6^) of cells is subcutaneously (s.c.) injected to the middle of the shaved area with a 291/2-G insulin syringe (EXEL). For tumor growth experiments, intravenous (i.v.) injection of DL80 (17 μg), hIL-2 (7.5 μg, PeproTech), TA99 (125 μg, BioXCell) and PBS were done on indicated days. For flow cytometry analysis, a single i.v. administration of DL80 (17 μg) or hIL-2 (7.5 μg) was done on day 12.

### Flow cytometry

The tumor was isolated from the mice (n = 5) 1 or 3 days after a single dose treatment. Cell suspensions were harvested by mechanical dissection and Collagenase IV Cocktail digestion (formulated as 7.4 kU Collagenase Type 4, 53 kU Deoxyribonuclease I, 20 mg Soybean Trypsin Inhibitor in 10 mL cocktail; Worthington Biochemical). The suspensions were passed through 40 μm cell strainers to ensure singularity. Cells were first treated with BioWhittaker ACK lysing buffer (Lonza), followed by TruStain FcX (BioLegend) and stained with antibodies against CD45 (clone 30-F11), CD3 (clone 17A2), CD8a (clone 53-6.7), NK1.1 (clone PK136). All antibodies are purchased from Biolegend. All samples were analyzed on a BD LSRFortessa cell analyzer. Data analysis was performed by Flowjo and Prism GraphPad.

## Acknowledgment

This work was supported by the Pitt SOM start-up funds (Y.S.), a UPMC Aging Institute pilot fund (Y.S.), and a grant from the Alzheimer’s Association and Michael J. Fox Foundation (Y.S.).

## Contributions

Y.S and Z.S conceived the research. Y.S. supervised the studies. Y.X and Y.S identified the Nbs. Z.S and Y.X performed major experiments and data analysis with help from Y.S, S.V, Z.X, A.V, C.J, G.C, B.H, G.C, Z. Sang, J.L and K.C. D.S and C.C performed structural modeling. Thank P. Duprex (Vaccine center, Pitt) for his assistance in the SPR experiments, D.Goebe, for critical reading of the manuscript. Y.S and Z.S drafted the manuscript with input from all the authors.

## Completing Interests

The University of Pittsburgh has filed a provisional patent encompassing the technologies described in this manuscript.

